# Different patterns of visual processing for shapes in V1 of mouse and monkey

**DOI:** 10.1101/2022.03.31.486346

**Authors:** Eve Ben Yehoshua, Hamutal Slovin

## Abstract

Recently, mice became a popular research model for the visual system including objects processing. However, while primates use their fovea to inspect objects at high resolution, mice have low visual acuity and lack a foveal region. Thus, how objects are encoded in the visual cortex of mice and whether monkeys and mice use similar neural mechanisms to discriminate between shapes, is unknown. Using voltage sensitive dye imaging, we investigated spatio-temporal patterns of population response in the primary visual cortex (V1) of mice and monkeys to shape contours. Neural responses in mice showed blurred spatial patterns, low correlation to the original stimuli and only subtle spatial differences between the shapes. In contrast, the neural responses in monkey’s V1 fovea, showed prominent differences between the shapes at sub-degree resolution with high resemblance to the original shapes, in agreement with a topographic code. Furthermore, the spatial patterns of neural synchronization in monkey’s V1 suggested contour binding by synchrony, but not in mice V1. Finally, monkey’s V1 responses at larger eccentricities, resembled more to those in mice. These results may suggest that to discriminate between shapes, mice rely on neural processing of low-level cues, whereas monkeys rely more on a topographic code.

## Introduction

How visual stimuli are processed and then transformed into behavioral actions is a key question in neuroscience. Past neurophysiological studies of cortical visual processing were done mostly in primates and carnivores which rely heavily on their visual system. However, recently rodents became a popular model in this field due to the wealth of experimental tools available for these animals. Neurophysiological studies reported that neurons in the primary visual cortex (V1) of mice and rats share many known hallmark features that were previously reported in primates. V1 neurons in mice and rats respond to contrast and luminance changes (Dai and Wang, 2012) and show tuning curves to orientation (Dräger, 1975; Niell and Stryker, 2008; Glickfeld, Histed and Maunsell, 2013) spatial and temporal frequencies (Marshel *et al*., 2011; Palagina, Meyer and Smirnakis, 2017; Marques *et al*., 2018). However, key functional maps such as ocular dominance or orientation columns, which are well established in higher mammals, were not found in the mouse visual cortex (Blasdel and Salama, 1986; Ts’o *et al*., 1990; Ohki and Reid, 2007; Van Hooser, 2007). Moreover, while primate use their fovea to inspect objects at high details, mice lack both fine-scale visual acuity and the specialized fovea region in the retina (Prusky, West and Douglas, 2000; Van Hooser, 2007; Huberman and Niell, 2011). Thus, it remains poorly understood how the visual system in rodents processes visual objects.

Rodents and non-human primates (NHP) show retinotopic organization of the visual space. The topography in V1 of mice is similar to that found in other mammals: the upper visual field (VF) is projected onto the posterior parts of V1 and the lower VF is projected to the anterior part of V1 (Schuett, Bonhoeffer and Hübener, 2002; Wang and Burkhalter, 2007; Ji *et al*., 2015a; Zhuang *et al*., 2017). However, mice have large receptive fields (RFs, ∼20 deg in V1) in contrast to the sub-degree RFs in monkeys’ foveal region (Métin, Godement and Imbert, 1988; Niell and Stryker, 2008). Due to the small RF size in monkeys’ fovea, the spatial patterns of neural activation evoked simple shape stimuli can show a topographic code, namely, they resembles roughly the shape of the original stimuli (Zurawel, Shamir and Slovin, 2016; Macknik *et al*., 2019).

The large visual RFs, lack of fovea and the poor spatial acuity put into question the ability of mice to discrimination between visual stimuli. Yet, behavioral studies showed that mice can discriminate between visual stimuli comprised of low level features such as contrast and orientation (Andermann, Kerlin and Reid, 2010; Busse *et al*., 2011; Lee *et al*., 2012). In addition, mice can perform higher visual functions such as figure-ground segregation (Schnabel *et al*., 2018; but see (Luongo *et al*., 2021) as well as natural image processing (Yu *et al*., 2018). Object recognition and discrimination were attributed mainly to humans and NHP, however, recent studies reported that mice can perform tasks involving object and shape recognition (Bussey, Saksida and Rothblat, 2001; Brigman *et al*., 2005). Yet it remains an open question what are the neuronal mechanisms involved in these higher visual functions.

While humans and monkeys have a formidable complex visual system that is considered to recognize objects holistically, rodents have a much simpler visual system. It remains controversial whether rodents can extract object-defining visual features in similar way as primates. It has been suggested that rodents use low-level cues (e.g., local luminance or orientation) to discriminate different shapes. Minini & Jeffery (2006) reported that rats relied on low-level cues (specifically luminance differences in the lower hemifield) to discriminate shapes (Minini and Jeffery, 2006). In contrast, other studies reported that rats can learn complex tasks of shape discrimination while exhibiting tolerance to object transformation such as changes in pose, illuminance, size, position etc. (Zoccolan *et al*., 2009; Tafazoli, di Filippo and Zoccolan, 2012; Vermaercke and Op De Beeck, 2012; Alemi-Neissi, Rosselli and Zoccolan, 2013; Vinken, Vermaercke and Op de Beeck, 2014; Tafazoli *et al*., 2017). This visual function was considered until recently to be a unique capability of the visual system in primates (Ratan Murty and Arun, 2017a).

In the present study, we investigated the neuronal mechanism in V1 involved in visual processing of shape contours in lightly anesthetized mice and in a fixating monkey. We used voltage sensitive dye imaging (VSDI) which enables to measure the spatio-temporal patterns evoked by shape stimuli, at high spatial (meso-scale) and temporal resolution (Shoham *et al*., 1999; Slovin *et al*., 2002). We found that V1 activity in both species can discriminate between different shapes but may use a different neural strategy. V1 activity in mice was characterized by a wide spread activation with peak response at the contour’s center with only small visible differences between shapes. In contrast, the evoked spatial patterns of neural activation in monkey’s V1 resembled the shape of the original stimuli with a clear hole at the center. The foveal representation in V1 of primates demonstrated clear neural differences at sub-degree resolution of the shapes. Furthermore, in order to test feature binding by synchrony (Gray, 1999; Singer, 1999), we computed synchrony maps using seed correlation. The pattern of neural synchronization suggested contour binding in monkey’s V1, but not in mice. Taken together, these findings reveal differences of neural responses for object discrimination in primates and mice, suggesting that mice may rely on local stimulus features i.e. low-level cues to discriminate between pair of shapes whereas primates rely more on a topographic approach.

## Materials and methods

### Mice data

#### Animals

A total of 25 adult mice male (C57BL\6JOlaHsd; 12-15 weeks) were used in this study. All experimental and surgical procedures were carried out according to the NIH guidelines, approved by the Animal Care and Use Guidelines Committee of Bar-Ilan University and supervised by the Israeli authorities for animal experiments.

#### Surgery

Craniotomy was performed under deep isoflurane anesthesia. The mouse was ventilated using a SomnoSuite Small Animal Anesthesia System facemask throughout the surgery and imaging. A chamber was cemented to the cranium with dental acrylic cement over the visual cortex in the left hemisphere (centered at ∼3 mm lateral from midline and ∼2.5 mm posterior from lambda). A 5 mm diameter craniotomy was performed and the dura mater was removed in order to expose an imaging window over the visual cortices. The cortical surface was then carefully washed with artificial cerebrospinal fluid (ACSF) and prepared for staining. The cortical staining lasted for 1.5-2 hr, the cortical surface was then re-washed with ACSF until the solution was clear. Next, the chamber was filled with transparent agar and covered with transparent Perspex. The mouse was placed on Harvard Small Animal Physiological Monitoring System (Harvard Apparatus) and was maintained on a body temperature at 37^0^C and oxygen saturation > 90% in room air. Before VSDI onset Carpofen (5 mg/kg s.c.) was given to the animals and 30 minutes before the VSDI onset animals were switched to light anesthesia (0.5% isoflurane).

#### Visual stimulation

Visual stimuli were displayed on a LCD screen (27×48 cm, 60 Hz refresh rate) placed ∼15 cm from the mouse. The stimuli were presented to the right eye (the left eye was covered with an opaque pad) while imaging was performed from the left hemisphere. We used three different sets of visual stimuli in different experiments (Fig. 1A):

1. Retinotopic stimuli for corner detection: mice (n=25) were presented with a small white square (85 cd/m^2^; square size: 5×5 deg; Fig. 1A top) that appeared on each trial in one out of six different positions in the VF. Together, the locations of the small white squares spanned the corners and the mid upper and lower edges of an imaginary large square contour (40×40 deg; Fig. 1B left insets). Stimuli were presented over a dark background (0.05 cd/m^2^) for 300 ms that was followed by inter-trial intervals of dark background.
2. Shape stimuli: animals were presented with four different white (85 cd/m^2^; 100% contrast) shape contours presented over a dark background (0.05 cd/m^2^) for 300 ms. The shapes included: square, triangle, inverted-triangle and circle (the circle and the triangles shapes were circumscribed in the square shape; see Fig. 1A middle and Fig. 2A left). Size: 40×40 deg; contour width: 5 deg; position: center of screen. On each trial, a randomly selected shape stimulus was presented followed by inter-trial intervals of dark background. The average number of trials for each stimulus was: square and circle: 19.6±5.85 (mean±1STD, 11 animals), triangle and inverted triangle: 20.3±5.3 (mean±1STD, 19 animals). To investigate the influence of contrast on shape processing, we used white shape stimuli at 50% contrast (background 42.5 cd/m^2^, n=6 animals) in a separate experiment.
3. Scrambled stimuli: to study local feature processing versus coherent shape processing we generated scrambled versions of the shape stimuli (Fig. 1A bottom). The scrambled stimuli preserved the local features of the original stimuli and the total luminance content while abolishing the coherent shape contour. For this purpose, the triangle and inverted-triangle images were divided into a matrix of 5×5 square segments, each segment 8×8 deg. We then randomly shuffled these segments (except for the vertices of the triangles) and created 10 different scrambled stimuli. The scrambled images were presented for 300 ms over a black background, n=8 animals.

**Figure 1.**
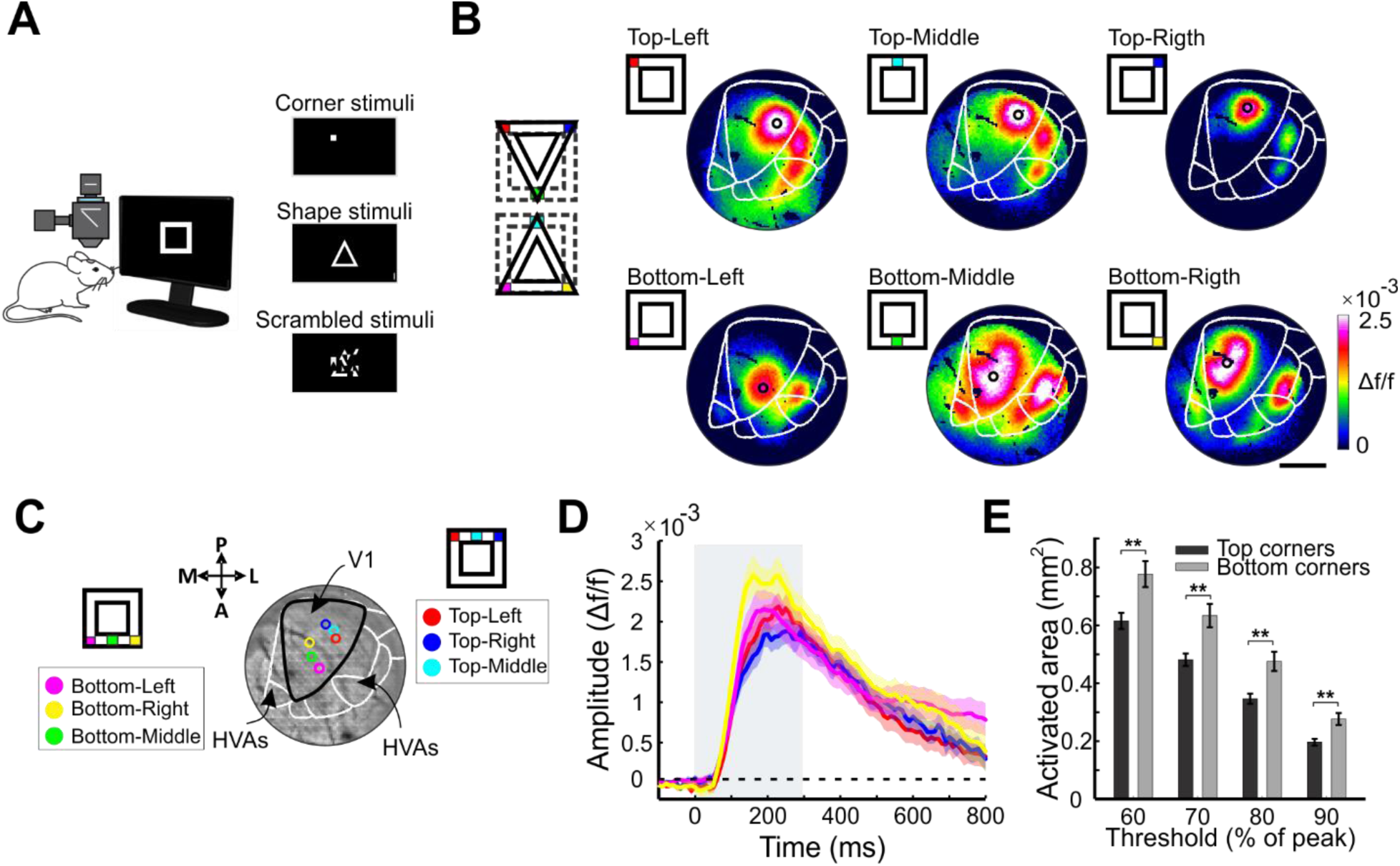
Experimental paradigm and retinotopic mapping of the shape edges and corners. (A) Left: schematic illustration of the VSDI experimental setup in anesthetized mice. Right: the visual stimuli (see Materials and Methods): corner stimuli (used for retinotopic mapping), shape contours and scrambled shape contours. (B) Retinotopic mapping of V1 using the corner stimuli, example session. Left: Illustration of the six corner stimuli and their location on the triangle and square shapes. Right: VSD maps at peak response amplitude (averaged 150-300 ms after stimulus onset) for each corner stimulus (see left insets). The fluorescence level (Δf/f) is color coded (here and in all other figures). The black circles depict the center of the corner ROIs for each stimulus (see Materials and Methods). The white contour depicts a template for visual cortical areas in the mouse (here and in all figures; see Materials and Methods). Scale bar is 1mm. (C) Corner ROIs for each corner stimulus superimposed on the image of the blood vessel pattern in V1. The color encodes the stimulus position on the shape contour. (D) Time course of evoked response for four corner stimuli in the square shape (color coded as in C; color shaded area represents ±1 SEM over animals, n=25). Gray rectangle represents the stimulus duration (here and in all other figures). (E) The size of activated area in V1, computed for increasing thresholds (60-90 % of peak response) for the top (black) and bottom (gray) corners stimuli. Wilcoxon rank-sum test: ** p<0.005.

**Figure 2.**
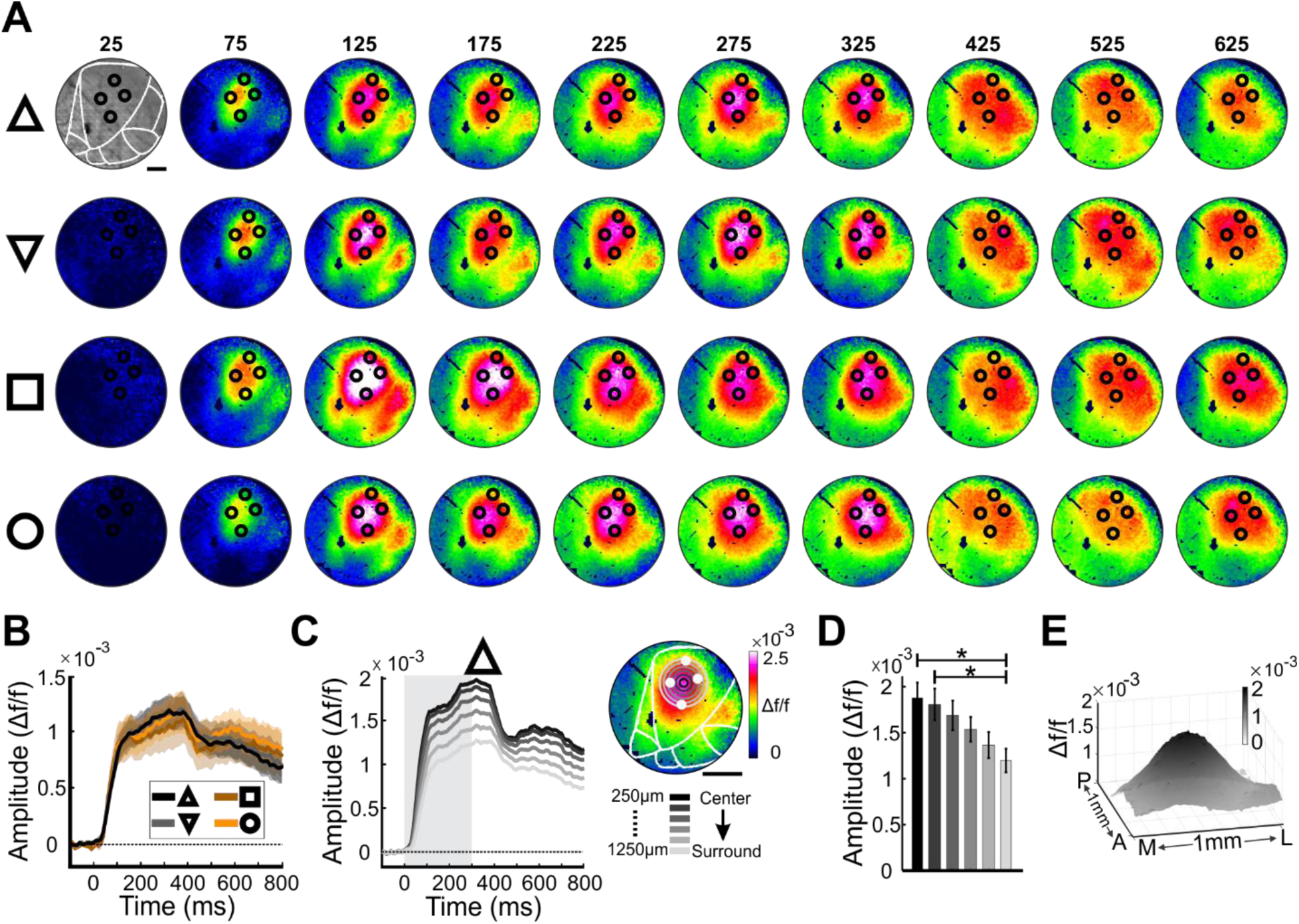
Spatio-temporal patterns of neuronal population evoked by shape contour stimuli. (A) A time sequence of VSD maps evoked by the various shape contours, an example session. The shapes are depicted on the left. Each map was averaged over a time window of 50 ms (the numbers above each map refers to the mid time window, following visual stimulus onset). Color bar is as in C. The map on top row, at t=25 shows the blood vessels pattern of the imaged window. The black circles denote the corner ROI location of the square shape. (B) Time course of the VSD signal in V1, for the shape stimuli, averaged across all animals (see Materials and Methods; square and circle, n=11; triangle and inverted-triangle, n=19). There are no-significant amplitude differences between the shapes (Kruskal-Wallis test, p=0.78±0.03, mean±1SEM across frames and shapes). (C) Space-time analysis using ring ROIs (see Materials and Methods). Right: Schematic illustration of ring ROIs superimposed on the VSD map evoked by the triangle shape. The distance between two successive rings is 200 µm. White filled circles denote the corners ROIs. Left: The VSD TC for the triangle shape average across all sessions (n=19), in the ring ROIs. The color of the curves denotes the ring distance, from the inner ring (black circle) to the outer ring (gray circles). (D) Peak response (time averaged 200-350 ms post stimulus onset) for all sessions in each ring ROI, same data as in C. Kruskal-Wallis test: * p=0.007 with *post hoc* Tukey tests: comparison of center ring with most two outer rings p<0.05). (E). 3D VSD activation map for the triangle stimulus, aligned on the shape center and averaged across all animals (triangle, n=19). Same data as in C.

#### Voltage-sensitive dye imaging

VSDI was performed in lightly anesthetized mice (0.5% isoflurane). We recently showed that V1 response in this sedated state shows only minor changes from the awake state (Nivnsky Margalit *et al*., 2022). We used the MicamUltima system with spatial resolution of 100×100 pixels/frame (imaged area of 5×5 mm²; thus, each pixel covers a cortical area of 50×50 μm²) and temporal resolution of 100 Hz (i.e. frame duration 10 ms). During imaging, the exposed cortex was illuminated using an epi-illumination stage with an appropriate excitation filter (peak transmission 630 nm, width at half height 10 nm) and a dichroic mirror (DRLP 650 nm), both from Omega Optical, Brattleboro, VT, USA. In order to collect the fluorescence and reject stray excitation light, barrier post-filter was placed above the diachronic mirror (RG 665 nm, Schott, Mainz, Germany).

#### Basic VSDI analysis

Data analysis was done using MATLAB software. The basic analysis of the VSD signal is detailed elsewhere (Ayzenshtat *et al*., 2010; Margalit and Slovin, 2018) . Briefly, to remove the background fluorescence levels, each pixel fluorescence was divided by its baseline fluorescence level (averaged over few frames before stimulation onset). The heart beat artifact and the photo bleaching effect were removed by subtracting the average blank trials response (no visual stimulation) from the stimulated trials. Thus, the VSD signal (Δf/f) reflects the changes in fluorescence relative to the blank condition. VSD signal and maps were computed by averaging over all trials.

#### Regions of interests (ROIs) analysis

Note, we used only V1 area for our analyses.

<u>Corner ROIs:</u> we defined a ROI for each of the corner stimuli in V1 using the following steps. First, for each corner stimulus, we computed a VSD map (average over 50-200 ms after stimulus onset) and selected a general large region around the activation patch in V1 and set a threshold of maximal response minus 1 STD (across pixels within this general region). Next, we defined a ROI that included all pixels with response amplitude larger than the threshold. Finally, we detected the center of this ROI and set a circle ROI comprised of 25 pixels (Fig. 1B black circles and Fig. 1C).

<u>Fitting a template of mouse visual areas:</u> we used V1 anatomical coordinates (see Surgery section) to identify the VSD response in V1. To determine the border of V1 area, we superimposed a template of mouse visual areas (adapted from (Marshel *et al*., 2011) on the VSD maps. We used a linear transformation (rotation and translation) to fit the template to the retinotopic activation patches in V1 and in HVAs. We verified that the visual stimuli in the upper and lower VF fell well within the retinotopic map of the upper and lower VF in V1 and in HVAs, respectively (for more information see supplementary Fig. S2 (Margalit *et al*., 2022).

<u>Computing the VSD response for the shape stimuli in V1:</u> to compute the time course (TC) of the VSD signal in V1 evoked by shape stimuli we performed the following steps: activation maps were averaged at 150-300 ms post stimulus onset. Next, we selected all pixels with response amplitude >= 70% of peak activation. The selected pixels were defined as the V1 ROI. To obtain the TC of VSD response for each shape, the VSD signal was averaged across all pixels within V1 ROI (Fig. 2B).

#### Similarity between VSD maps of shape stimuli

To evaluate the similarity between activation patterns of any two shapes (shape-pair), we calculated the pixel-wise Pearson correlation coefficient (r) for each shape-pair maps in V1 area. VSD maps were averaged at peak response (150-300 ms post stimulus onset, see Fig. 6A right) or during baseline (150-50 ms before stimulus onset). Finally, correlation values were averaged across sessions.

#### Ring ROI analysis for shape stimuli

Ring ROI analysis was applied to quantify and compare the VSD response amplitude from center of activation outwards, i.e. towards more peripheral regions (within V1 area). In mice we generated a continuous set of non-overlapping circle rings, aligned over the center of the 4 corner ROIs (i.e. the square center). The rings were separated by 200µm from each other (see schematic illustration in Fig. 2C right and Fig. S1B).

#### Computing the point-spread function (PSF)

The small retinotopic stimuli (i.e. corner stimuli) evoked a population response with a large spatial spread in V1 (Fig. 1B). For each corner stimulus, we computed the averaged VSD map after stimulus onset (150–300 ms). Next, this map was normalized to peak response, and a threshold of 40% from peak response was set (Fig. 3Ai, black contour). Next, a 2D Gaussian was fitted to the threshold map (Fig. 3Aii) and was defined as the PSF.

#### PSF convolution and computing the expected pattern

In order to compute the expected pattern of activation for the shape stimuli, the stimulus projection on V1 was convolved with a PSF. The projection of the shape contour from the VF to V1 was done by linearly connected the corner-ROIs on V1 (Fig. 3Aiii and see ROI analysis in Materials and Methods). To obtain the expected pattern of activation in V1, we used two approaches: (1) the stimulus projection was convolved with a single PSF (Fig. 3A) (2) each corner (and neighboring edges) of the projected stimulus was convolved with the PSF that was computed from the corner-ROI of that corner (for example, top middle corner with the upper part of the triangle). Finally, the convolutions of all segments were averaged to obtain the expected pattern for the entire stimulus. Using the latter approach, we took into the consideration that PSF of the lower VF had a larger spatial spread than in the upper VF (Fig. 1E). To evaluate the similarity between the observed and predicted activation patterns for each stimulus shape, we defined a ROI of 40% (Fig. 3B top, red contour) from peak response for observed map. Next, we calculated the Pearson correlation coefficient (r) between the measured and predicted patterns of the same shape for pixels in the ROI (Fig. 3B).

**Figure 3.**
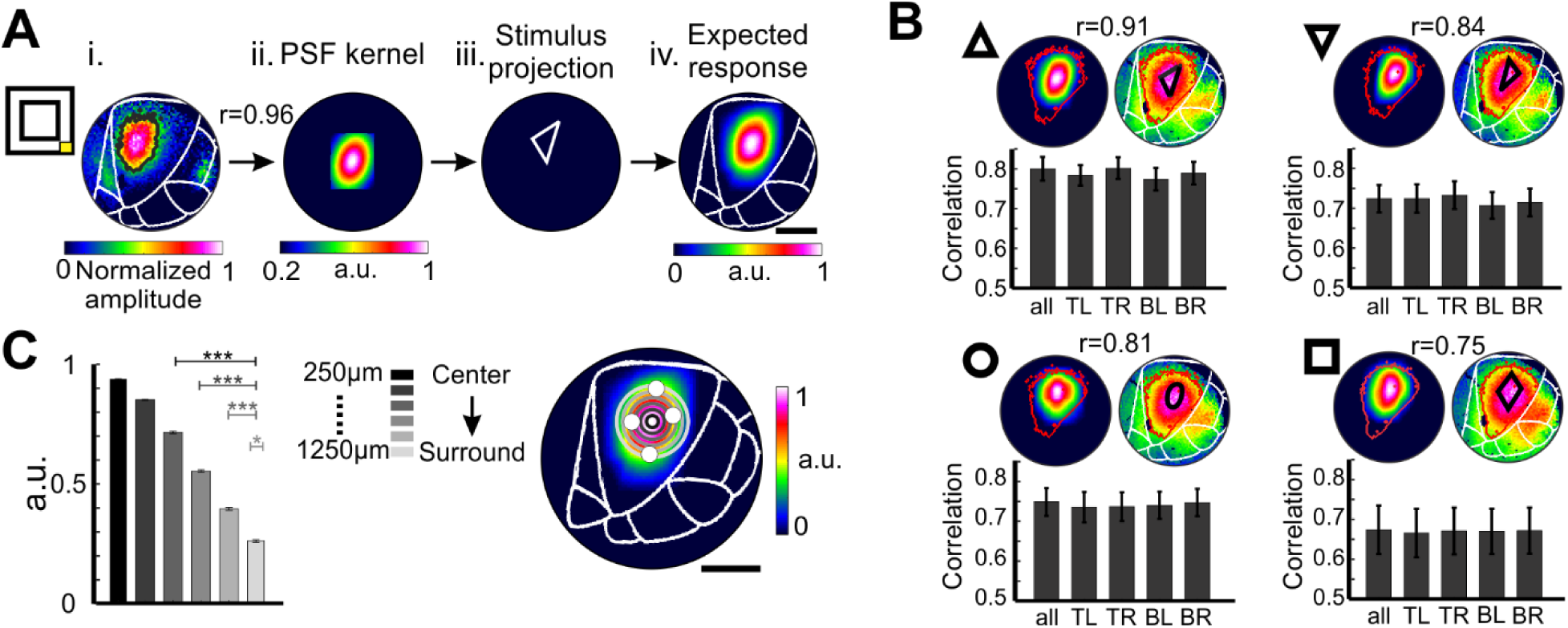
Computing the predicted response for the shape using the point-spread function and model performance. (A) Point-spread function fitted for a corner stimulus and predicted response for the triangle shape. From left to right: (i) The normalized VSD map evoked by the corner stimulus. The black contour depicts the pixels above threshold of 40% of peak response. The VSD map was averaged over 150-300ms post stimulus onset. (ii) The PSF kernel: a 2D Gaussian model was fitted for the response pattern (Pearson correlation coefficient, r=0.96; see Materials and Methods). (iii) The stimulus projection of the triangle’s contour on V1 (based on retinotopic mapping of the triangle corners). (iv) The expected response pattern computed from the convolution of the PSF kernel with the triangle projection on V1. (B) Correlation between observed and predicted maps. Top: example of measured and predicted VSD map for each stimulus shape (r between the maps is depicted above). Red contour depicts the pixels used for correlation (corresponding to pixels above 40% of peak VSD response). Bottom: grand analysis of the correlation between the observed and predicted maps for each type of stimulus, averaged across animals (triangle and inverted-triangle, n=19; rectangle and circle, n=11. Correlations were computed for the different PSFs estimation: top-left (TL), top-right (TR), bottom-left (BL) and bottom-right (BR) corners. ‘all’ represents the correlation when the PSF from each corner was convolved with the corresponding part of the stimulus (see Materials and Methods). (C) Ring analysis as in Fig. 2C-D but for the predicted response of triangle stimulus (Kruskal-Wallis test: p<0.0001 with *post hoc* Tukey tests: center ring with ring at 650 µm and larger p<0.01; ring at 450 µm with rings at 850 µm and larger p<0.01; ring at 650 µm with rings at 1050 µm and larger p<0.01; ring at 850 µm with ring at 1250 µm p<0.05.

#### Spatial cuts, spatial profile and space-time maps for shape stimuli

To quantify the spatial extent of the visually evoked activity in V1, we computed a spatial cut on the VSD maps at 3 locations along the shape maps (roughly along the posterior-anterior axis; Fig. 4A). The position of each spatial profile was aligned on the two top corner ROIs (posterior cut; Fig. 4A top right), two bottom ROIs (anterior cut; Fig. 4A top left) and at the center ROI (middle cut; Fig. 4A top middle). Next, we averaged the VSD signal across the width of the spatial cut to obtain spatial profile and compute the space-time plots (Fig. 4A bottom). Finally, we computed the width of the spatial profile (Fig. 4B) using the following steps: (i) averaged the space-time map over 150-300 ms. (ii) set a threshold of 80% from peak response and found the most outer pixels on each side of the spatial profile crossing the threshold. (iii) used a linear interpolation to find the accurate location of crossing the threshold. This approach enabled us to estimate the width of the spatial profile at higher accuracy than the imaged pixel size (50 μm).

**Figure 4.**
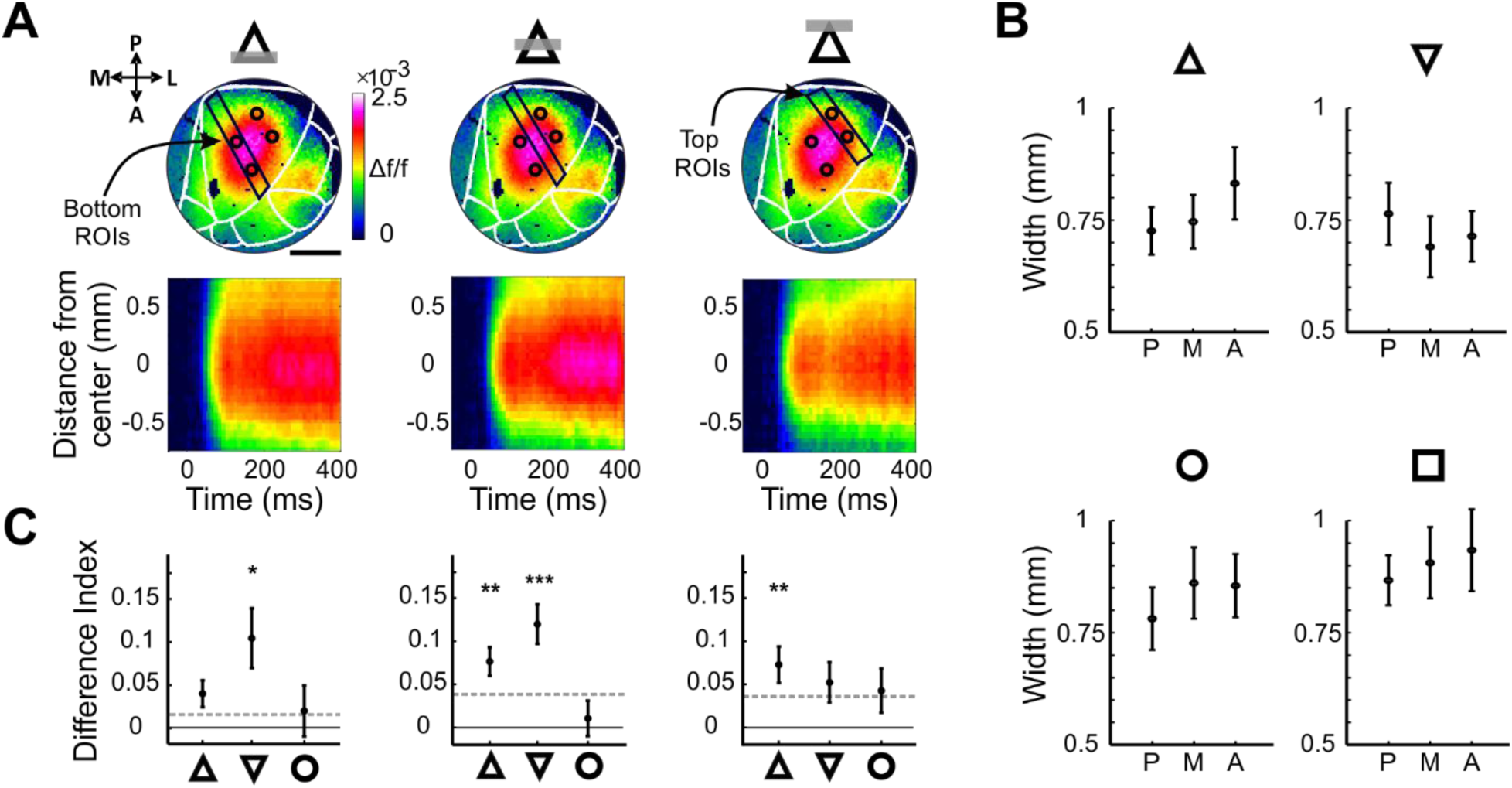
Narrow and wide parts of the shape contour generate wide and narrow spatial profile of cortical response. (A) Top: The spatial cuts that were used for space-time analysis are depicted as black rectangle superimposed on the VSD map. The spatial cuts were positioned roughly along the anterior-posterior axis in V1, from left to right: Anterior: positioned over the two anterior corner ROIs (corresponding to the lower edge of each stimulus). Middle: set to the center of activation (in between the four corner ROIs). Posterior: positioned over the two posterior corner ROIs (corresponding to the upper edge of each stimulus; see Martials and Methods). Bottom: Space vs. time maps computed from the spatial cuts in the top panels. The maps show the VSD response at increasing distances from the center of activation (distance=0; y-axis) as function of time (x-axis). (B) Width of the spatial profiles. Spatial profiles were computed from the spatial cuts (see Materials and Methods). Each panel depicts the width (in mm) of the 3 spatial profiles (A-anterior, M-middle, P-posterior), averaged over 150ms time-window, for each shape. The width was computed at 80% of peak response. (C) The width difference index (see Materials and Methods) between each shape and the square. The width differences index was calculated for the 3 spatial cuts (as in A). Wilcoxon sign-rank test from zero: * p<0.05, ** p<0.01, *** p<0.001. Dashed-line represented the expected threshold (see Materials and Methods).

To quantify the differences of spatial profile width between shape stimuli and to normalize to the PSF size in V1, we defined a Difference index measure (Equation 1; Fig. 4C):

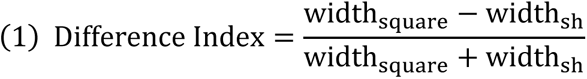

Where width_sh_ are the width of spatial profile (one of the 3 locations) for the triangles and circle shapes and width_square_ are the width of square shape in the same spatial profile location. We used the width of the square as a normalized measure because of its symmetrical shape. Positive difference index values mean larger width for the square stimulus, zero values mean no difference between the shape and the square and negative values mean larger width values for the shape stimulus. The difference index was calculated separately for each session and shape-square pair. To evaluate the statistical significance, we computed a shuffled difference index, for each spatial cut. We calculated the difference index between one random square (for all sessions) and a random shape. This was repeated over 100 permutations. Threshold was set as the averaged across permutations of the shuffled difference index (dash-line Fig. 4C).

#### *STD maps* for pairs of shape stimuli

To identify the pixels that are informative to discriminate between pair of shapes we performed the following steps, for each session: (i) activation maps were averaged at 150-300 ms post stimulus onset. Then we compute a differential map by subtracting one map from the other. (ii) Next, we computed mean and STD value in the differential map, across all pixels in V1. (iii) We then set two thresholds of Mean ±1 STD and defined three classes of pixels. Pixels with values <= Mean-1STD, depicted by brown color (represented by -1), pixels with values >=Mean+1 STD, depicted by yellow color (represented by +1), pixels with values within Mean±1 STD depicted by orange color (represented by 0). Examples of the resulted maps appear in Figure 5A. Pixels with values larger than ±1 STD denote pixels that carry information for shape discrimination. (iv) Finally, we computed the center of mass separately for the two classes of pixels (blue dots in Fig. 5A). To find the center-of-mass we used unsupervised General Mixture model (GMM). Note, we repeated this analysis on baseline time-window (Fig. 5A).

**Figure 5.**
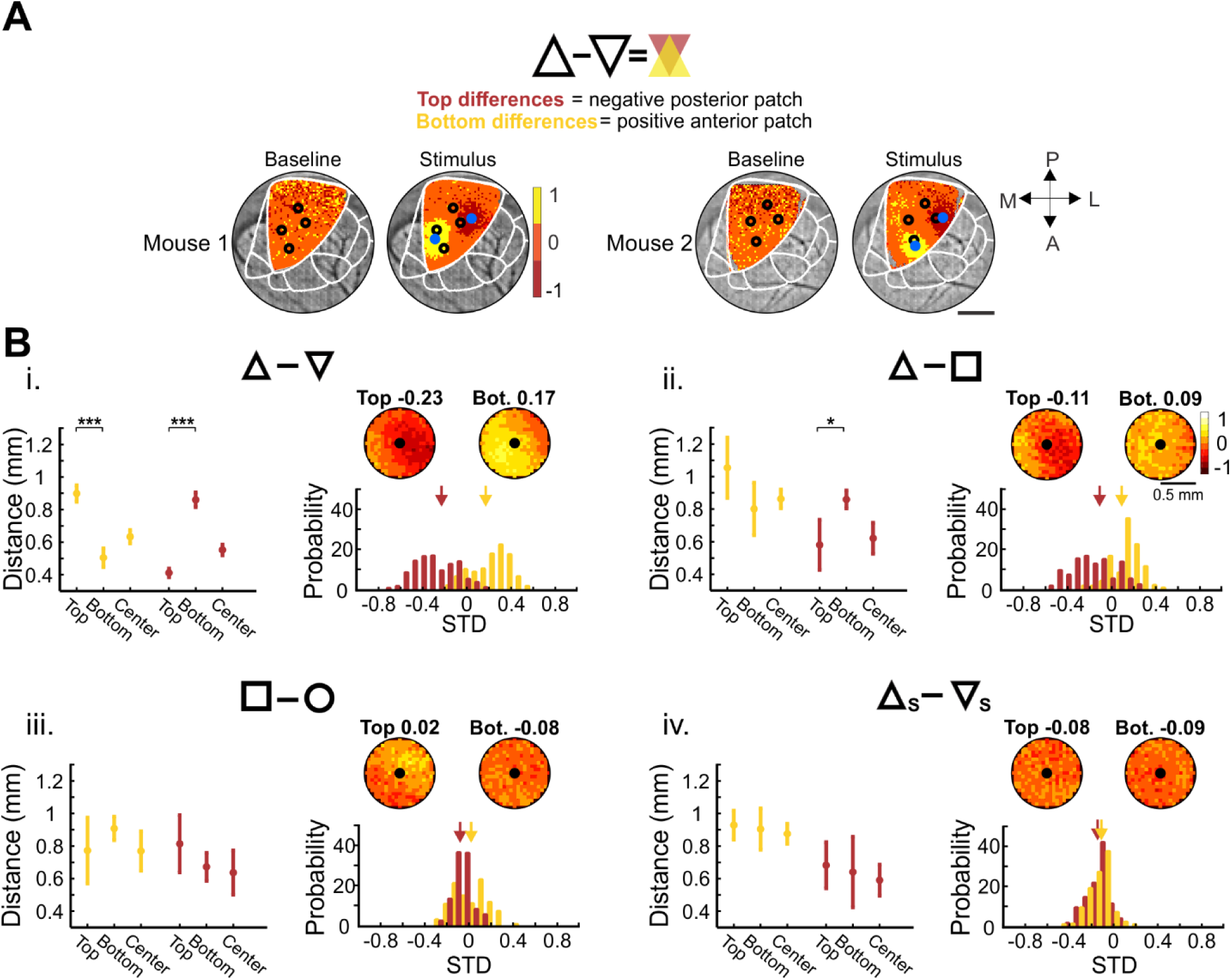
STD maps for the activation maps show patches at non-overlapping regions of the stimuli pairs. **(A)** Example of STD maps for a triangles-pair (triangle and inverted-triangle). The maps were calculated by computing the response difference between the two stimuli evoked maps i.e. a differential map. The yellow and red pixels denote pixels with response difference crossing a threshold of +1STD (positive; yellow) or -1STD (negative; brown) from the mean value of the differential map (see Materials and Methods). For each example animal, the left and right maps represent STD maps at baseline activity and stimulus time-window, respectively, with pixels color encoded relative to threshold. Blue dot depicts the center-of-mass for the negative or positive patches (see Materials and Methods). **(B)** i: Left: The Euclidean distance (mm) between the center-of-mass location for positive (yellow) and negative (brown) patches to the top, bottom corner ROIs or center ROI in triangle pair. Right bottom: The STD distribution around the top and bottom ROI, for the triangle pair. Right top: STD Maps averaged for top and bottom corners ROIs, across all animals (n=19, map radius is 0.5 mm). The number is the mean STD value in the map. ii-iv: same as in i, but for other stimulus-pairs (triangle-square and square-circle: n=11; scrambled triangle pair: n=8). Wilcoxon rank-sum tests: * p<0.05, *** p<0.001.

To study where the positive and negative patches within V1 are located, we calculated the distance between the patch center of mass to all 4 corner and center ROIs (Fig. 5B left panels). The distance to the two top corners or the two bottom corners showed similar results and thus were averaged.

Seed correlation analysis and synchronization maps in mice are described under Monkey data (see below).

### Monkey data

#### Animal, surgical procedures VSDI and dataset

One adult male monkey (Macaca fascicularis, 10 Kg) was used for the current study.

All methods were approved by the Animal Care and Use Guidelines Committee of Bar-Ilan University, supervised by the Israeli authorities for animal experiments and conformed to the NIH guidelines. The surgical, staining and VSD imaging procedure have been reported in detail elsewhere (Slovin *et al*., 2002). Briefly, a chamber was positioned over V1 area and VSDI was obtained from the left hemisphere. For VSDI we used the MicamUltima system with spatial resolution of 100×100 pixels/frame (imaged area of 17×17 mm²; thus, each pixel covers a cortical area of 170×170 μm²) and temporal resolution of 100 Hz (i.e. frame duration 10 ms). Dataset included: 9, 15 and 14 sessions for each stimulus location in the VF (Fig. 7) and 9 sessions were used for ring analysis (Fig. 6B-D). Correlation analysis was done on a total of 5 sessions: 3 circle sessions and 2 sessions for triangle and rectangle (Fig. 8Eii, 8F). A small part of the data (3 sessions) were used in a previous paper (Macknik *et al*., 2019), and here they were re-analyzed differently for a comparison with mice data.

**Figure 6.**
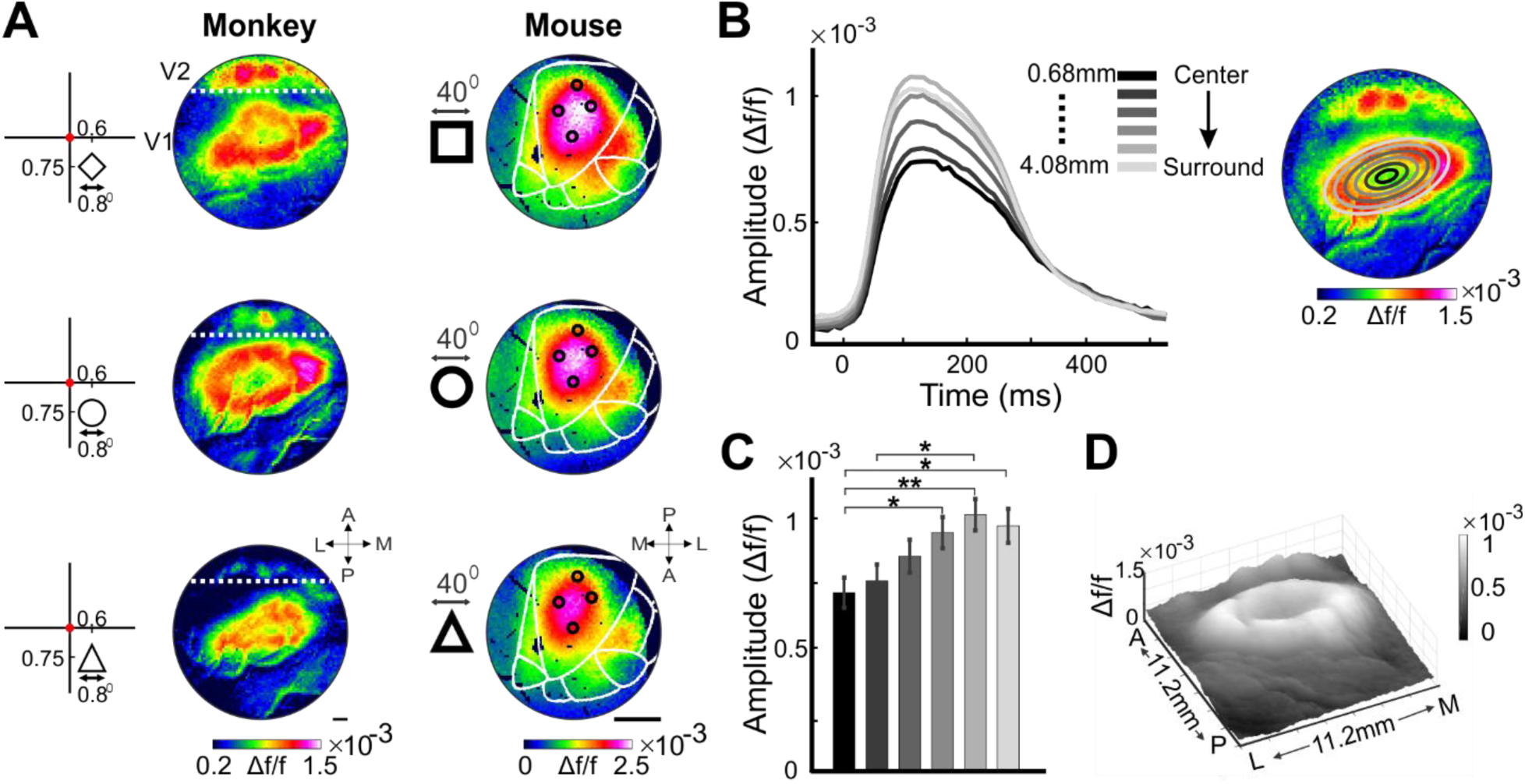
Comparison of shape processing in monkey’s V1 foveal representation and mouse V1. (A) VSD maps evoked by fine shape contours in V1 of a fixating monkey (left) and in V1 of a mouse (right). Left: shape contours stimuli (diameter=0.8°; contour width: 0.05°; positioned at 0.6° deg right to the vertical meridian and 0.75° below the horizontal meridian) were presented on a gray background while the animal was maintaining fixation on a small fixation point (red point, for illustration). Middle: VSD map at peak response for the shape stimuli. Data are from monkeys V1. Right: VSD maps at peak response for each stimulus (left inset). Same data as in Fig. 2A). Data are from mouse V1. (B) Time-course of the VSD signal evoked by a circle shape, in monkey, averaged across all sessions (n=9). The TC was computed for elliptical ROIs from the center of activation pattern (black ellipse) to the surround (gray ellipses). Gray scale indicates the distance (mm) from center, as presented in the right image. (C) Grand analysis of VSD peak response for all circle sessions (n=9), for each elliptical ROI. Response was averaged over a 100 ms window (130-230 ms post stimulus onset). Kruskal-Wallis test: p=0.0005 with *post hoc* Tukey tests: center ring with rings distance 2.04 mm and larger p<0.05; ring 1.36 mm with ring 4.08 mm p=0.02) (D) A 3D VSD map for the circle shape, same data as in B.

**Figure 7.**
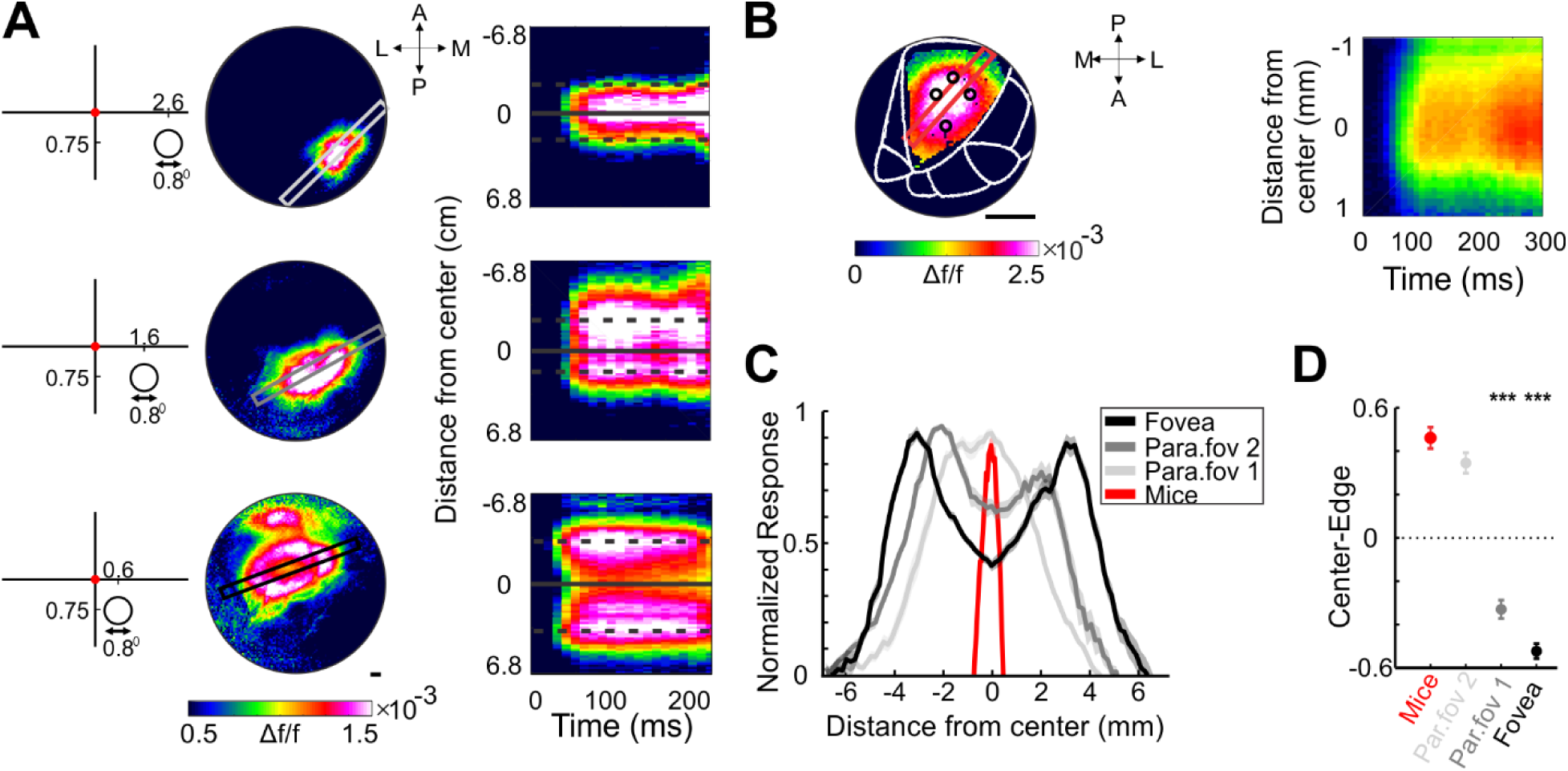
The VSD response pattern in monkey’s parafoveal area is more similar to the VSD response pattern in mouse V1. (A) VSD maps in monkey’s V1 evoked by a circle stimulus (0.8°) positioned at three different locations in the VF. Left: VSD peak response maps for each location of the circle. Top: stimulus coordinates 2.6° right to the VM and 0.75° below the HM; middle: 1.6°, 0.75°; bottom: 0.6°, 0.75°. Superimposed on the maps are spatial cuts (black and grays rectangles) that were used for computing the space-time maps (right). Right: Space-time maps computed from the spatial cuts in the left panels (see Materials and Methods). The maps show the VSD response at increasing distances from the center of activation (distance=0; y-axis) as function of time (x-axis). The center position of the circle is marked by a continuous horizontal line and the circle edges are marked by horizontal dashed lines. (B) Left: Peak response the VSD map for a circle stimulus (diameter=40°; contour width: 5°) in mouse V1. Superimposed on the map is a spatial cut (red rectangle) that was used for computing the space-time map. The spatial cut was positioned along the long-axis of the cortical activation. Right: Space-time map computed from the spatial cut in the left panel. (C) The spatial profile of the population response in A (black and gray curves) and in B (red curve), at peak response, aligned on the center of the response. Response was averaged over sessions (n=11 for mice and n=9,15, 14 for monkey’s fovea, parafovea-1 and parafovea-2, respectively). (D) Center to edge difference: the response difference between center and edge in calculated in mice (red symbol) and monkey (black and gray symbols) spatial profiles (from C). Wilcoxon rank-sum tests: ** p<0.01 for comparison between mice and monkey.

**Figure 8.**
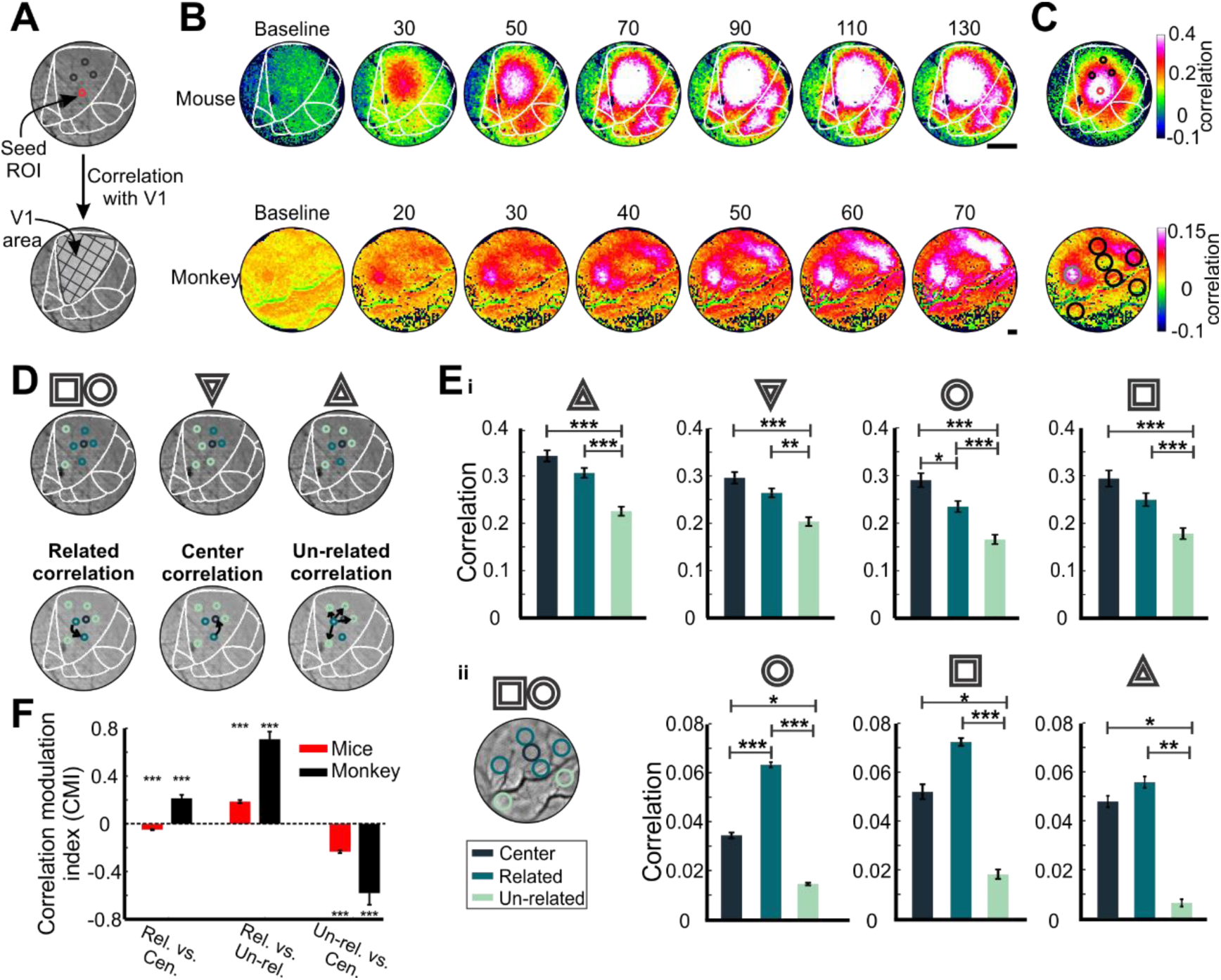
Seed correlation maps in mice and monkey reveal the stimulus mapping in V1. (A) Schematic illustration of seed correlation analysis. The red circle on the top maps depicts the seed pixels. We computed the correlation of the seed pixels (corner ROIs) with all the pixels in V1 denoted by grid area at the bottom. (B) A time sequence of correlation maps in mouse V1 (top row, circle stimulus) and in monkey (bottom row, circle stimulus). Each map was averaged over a time window of 100 ms (the numbers above the maps depict the mid time window, after visual stimulus onset). (C) A correlation map in mice and monkey for a circle shape. The seed ROIs (red in mouse, gray in monkey) is located on the contour. The correlation maps were averaged at 50-150 ms after stimuli onset for mice and 50-100 ms for monkey. (D) Top: The 3 types of ROIs (color coding of the ROIs appears in Eii) defined for each shape: *(1) Related ROIs*. *(2) Un-related ROIs*. *(3) Center ROI*. Bottom: Schematic illustration of the 3 correlation types. *(1) Related correlation*: the correlation between a seed related ROI to all other related ROIs; *(2) Un-related correlation*: the correlation between seed related ROI to all other un-related ROIs. *(3) Center correlation*: the correlation between the center ROI to all seed related ROIs. (E) Grand analysis of the correlation in mice (i) and monkey (ii). Mice (i): correlation values were averaged over a time window (50-150ms post stimulus onset), and across sessions (n=11 for rectangle and circle; n=19 for triangle and inverted-triangle). The colors indicate related, un-related and center correlations. Two-sided Wilcoxon rank-sum test: * p<0.05, ** p<0.01, *** p<0.001. Monkey (ii) Left: illustration of the 3 types of ROIs in monkey for rectangle and circle shapes. Right: same as (i) for monkey. Time window: 50-100ms post stimulus onset. (F) Correlation modulation index in mice (red) and monkey (black). The Δ correlation between the 3 correlation types as indicated below each bar. Wilcoxon sign-rank tests: p<0.001 for all bars.

#### Behavioral paradigm

The monkey was trained on a simple fixation task. On each trial, after a random fixation period (3-4 s) the monkey was presented with a briefly (300 ms) single visual stimulus as described below. The monkey was required to maintain tight fixation throughout the whole trial and was rewarded with a drop of juice for each correct trial. Stimulated trials were interleaved with blank trials, in which the monkey fixated but no visual stimulus appeared.

#### Experimental setup, visual stimuli and monitoring eye position

The experimental setup has been described in detail elsewhere (Ayzenshtat *et al*., 2012; Zurawel *et al*., 2014). Visual stimuli consisted of a black circle (dimeter=0.8 deg) square or triangle contours (Fig. 6A left). Stimulus contrast was 50% (gray background) and stimulus position was set to 0.6 deg, 1.6 deg, 2.6 deg, right to the VM and 0.75 deg below the horizontal meridian. Stimuli position were termed as foveal, parafoveal-1 and parafoveal-2 accordingly (Fig. 7A left). Contour width was 0.05 deg.

Eye positions were monitored using a monocular infrared eye tracker (Dr. Bouis, Karlsruhe, Germany) which resolves 0.1 deg, sampled at 1 kHz and recorded at 250 Hz. Only trials where the animal maintained a tight fixation were analyzed.

#### Ring ROI analysis

Ring ROI analysis was applied in order to: (i) quantify the VSD response amplitude from center of activation outwards (ii) compare the activation patterns in the monkey (Fig. 6B-C) to mice (Fig. 2C-D). For this purpose, we generated elliptical rings to fit the ROIs to the pattern of the response (see schematic illustration in Fig. 6B). The size of the minor and major axes of the ellipses increased at steps of 2-4 pixels.

#### Space-time maps: comparison between mice and monkey

To compare the activation patterns in mice and monkey, we used spatial cuts superimposed on the VSD maps in both species. In the monkey, we set the spatial cuts over the VSD maps evoked by a circle contour that appeared at different positions in the VF. Parafoveal-2: 2.6 deg right to the VM and 0.75deg below the HM (Fig. 7A top); Parafoveal-1: 1.6 deg, 0.75 deg (Fig. 7A middle); Foveal: 0.6 deg, 0.75 deg (Fig. 7A bottom). The spatial cuts were positioned along the long-axis of the VSD activation pattern. In mice we set a spatial cut along the elongated anterior-posterior axis of activation (positioned at the center of the 4 corners; Fig. 7B). Next, we averaged the VSD signal across the width of the spatial cuts to obtain the spatial profile and compute the space-time maps (monkey: Fig. 7A right; mice: Fig. 7B right). The normalized spatial profiles (normalized to peak response amplitude at 100-200 ms and 150-300 ms for monkey and mice, respectively) were averaged across sessions in the monkey (9, 15 and 14 sessions for fovea, parafovea-1 and parafovea-2 respectively) and over animals in mice (n=11; Fig. 7C). Finally, we calculated the center-edge index that is the response amplitude difference between the center and the edge (Fig. 7D). Positive center-edge values mean higher activation in center than edges, negative values mean higher activation in the edges and zero values mean similar activation at center and edges.

For each stimulus position, we defined the location of the center and edges. In the fovea and parafovea-1 positions, the center of the spatial profile (continuous horizontal line, Fig. 7A) was defined as the low activation region of the evoked response. The edges (dash horizontal line, Fig. 7A) were defined as peak activation regions in the spatial profile. In the parafovea-2 position and in mice, where there is one peak in the center, the center is defined as the high activation region. The edges in in mice were defined based on the linear interpolation between corner ROIs.

To define the stimulus edges on the cortical evoked response for the parafoveal-2 location in th monkey, a latency map was computed for each session. First, we selected pixels with significant evoked response to the visual stimulus (i.e. a minimal threshold of mean VSD response plus 3 STDs relative to baseline activity). Next, we selected pixels with VSD signal crossing this threshold for at least 4 consecutive time points. Then, a linear interpolation was used on the VSD signal to define the exact time for crossing the threshold. This time value for each pixel defined the latency map. Next, we defined a latency ROI that included all the pixels with latency values that are smaller than the median latency (i.e. pixels with shortest latency). We then set a spatial cut on the latency map and stimulus edges were defined as the border of the latency ROI.

#### Seed correlation analysis and synchronization maps in mice and monkey

To investigate the synchrony in V1 area (in mice or monkey), we computed the synchrony between neural populations by calculating the correlation at the single trial level. We first subtracted the mean evoked VSD time course (averaged across trials) from the time course of each pixel (Eq. 3; Brody, 1999). This was done separately for each stimulated condition. Using this procedure, we removed most of the neural activity that is directly related to the visual stimulus onset.

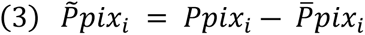

Where P_pix i_ is the population response in pixel i from a single trial and ^̄^*pix i* is the mean population response in pixel i across trials (we excluded from the mean response the specific trial). Using a sliding rectangular window of 10 frames (100 ms) we calculated the Pearson correlation coefficient (r; Eq. 4) between each pixel in a seed ROI (mice: 5×5 pixels, 250×250 μm^2^; monkey: 7×7 pixels, 1190×1190μm^2^) to all other pixels in the imaged window (Fig. 8A). The seed ROI was located at the spatial location of the corner ROIs.

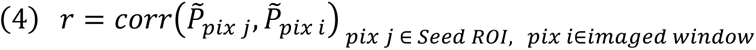

Where corr is Pearson correlation coefficient, 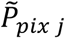 is the VSD signal of pixel j within the seed ROI and 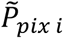 is the VSD response of pixel i in the imaged area. This was computed for each time window width (100 ms) along the time axis: -200ms to 450ms post stimulus onset for mice and -70 ms to 430 ms for monkey post stimulus onset, for each single trial. Next, single trial correlation maps were averaged, for the same seed ROI. Finally, synchrony map was defined as the mean r values between the seed ROI pixels to all other pixels in the imaged area over a time window (50-150 ms for mice and 50-100 for monkey post stimulus onset). The outcome of this procedure was a separate synchronization map, for each seed ROI (Fig. 8C). These correlation maps preserved the anatomical relation between the different areas and revealed the functional correlation between the seed ROI to V1 (correlation between seed pixels to themselves were excluded from the data).

To evaluate the statistical significance of the seed correlation maps we computed the shuffled correlations maps. We calculated the correlation (r) between the seed pixel in each trial to all other pixels in a randomly chosen trial of the same condition. This was repeated over 100 permutations. Next, we set a threshold for correlation (r values were averaged across windows located 50-150 ms and 50-100 ms post-stimulus onset for mice and monkey, respectively) as larger than the maximal value and lower than the minimal value of the averaged shuffled maps (averaged across permutations). Thus, in the real data, only pixels with correlation values exceeding this threshold were further analyzed.

Next, three types of ROIs were defined: (1) *Related ROIs*: shared corner ROIs i.e., ROI with overlapping visual input with the shape (for example bottom right, bottom left and top middle in triangle shape). (2) *Un-related ROIs*: corner ROIs with non-overlapping visual input for the shape (for example top left and top right in triangle shape). (3) *Center ROI*: located at the center of the shape contour, i.e., at the center of the 4 rectangle ROIs corners (see Fig. 8D top). We then calculated 3 types of correlations (Fig. 8D bottom): (1) *Related correlation*: the correlation between a seed related ROI to all other related ROIs. *(2) Un-related correlation*: the correlation between seed related ROI to all other un-related ROIs. (3) *Center correlation*: the correlation between all seed related ROIs with center ROI.

Finally, we computed Correlation-Modulation-index (CMI; Eq.5):

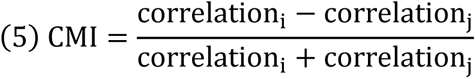

Where correlation_i_ is the related or un-related correlation and correlation_j_ is the un-related or center correlation. CM index was calculated separately for each session and for each corner ROI (Fig. 8F).

#### Statistical analysis

To compare the VSD response across different shape and between mice and monkey, nonparametric statistical tests were used. Signed-rank test to compare a population’s median to zero (Figs. 4C, 8F). To compare between medians of two populations we used Wilcoxon Rank-Sum test (Figs. 1E, 5B, 7D and 8E). For multiple-sample statistics we used Kruskal-Wallis test (with *post hoc* Tukey test; Figs. 2D, C3 and 6C). Data are presented as mean ± SEM, unless specified otherwise.

## Results

Mice are an important animal model to study the visual system, thus a main goal of this work was to investigate the neural processing of shape contours in their visual cortex. We first mapped the shapes onto the retinotopic map of V1 and then characterized the neuronal responses to the various shape contours and quantified their differences. While humans and NHPs use their fovea to inspect objects with high resolution, mice lack both fine-scale visual acuity and the specialized fovea region in the retina (Huberman and Niell, 2011). Thus, our next step was to compare shape processing in V1 of mice and monkeys while focusing on the foveal representation of the monkey’s V1. Cortical activity was measured using VSDI from the visual cortex of both mice and monkey (Fig. 1A and see Materials and Methods). This technique measures the membrane potential of neuronal populations from the cortex (mainly mid-to-upper layers) at high spatial (mesoscale, 50^2^ μm^2^/pixel) and temporal resolution (100 Hz) simultaneously (Shoham *et al*., 1999; Grinvald and Hildesheim, 2004; Berger *et al*., 2007). A main advantage of this technique is the combination of wide field imaging with high spatio-temporal resolution, which enables to visualize the complete cortical patterns evoked by the shape contour.

### Mapping the corners of shapes contour onto V1

To map the corners of the shape contour onto mouse V1, we used a set of small white squares (i.e., corner stimuli; Fig. 1A top) positioned at the corners and mid edges coordinates of the square and triangles shapes (Fig. 1B, left insets). Figure 1B right shows the VSD maps evoked by the corner stimuli in one animal. The VSD maps show clear patches of activation in V1 area as well as in higher visual areas (HVA; V1 and HVA are marked in Fig. 1C; this work focuses on V1 analysis, thus HVA were not further analyzed). As expected, the corner stimuli in the upper (Fig. 1B top) and lower (Fig. 1B bottom) VF evoked activation patches in the posterior and anterior parts of V1, accordingly. These findings are in accordance with the known VF mapping in mice V1 (Schuett, Bonhoeffer and Hübener, 2002; Wang and Burkhalter, 2007; Ji *et al*., 2015b; Zhuang *et al*., 2017). Next, for each corner stimulus, we defined a corner ROI and the center of this ROI is marked as a black circle on the VSD maps (Fig. 1B; see also Materials and Methods). Figure 1C shows the superposition of corner ROIs over the blood vessels map. Figure 1D depicts the grand analysis (averaged across all animals) time-course (TC) of the evoked population response for the corner stimuli of the square, demonstrating similar VSD response amplitude and dynamics (Kruskal-Wallis test, p=0.19±0.02, mean±1SEM over p values for all stimuli and frames).

Interestingly, the corner stimuli in the lower VF (mapped to the anterior part in V1) evoked activation patches that spread over a larger area as compared with the corner stimuli in the upper VF (Fig. 1B). The size of activated area in V1 was computed for different percentile thresholds for each corner stimulus (60%-90% of peak response). Figure 1E shows that the top corners evoked significantly smaller responses (in the posterior part of V1) than the responses evoked by bottom corners (in the anterior part of V1) for all percentile thresholds (mean±1SEM mm^2^, n=25; 60%: 0.62±0.03, 0.79±0.05 for top and bottom respectively; 70%: 0.48±0.02, 0.64±0.04 for top and bottom respectively; 80%: 0.35±0.02, 0.48±0.03 for top and bottom respectively; 90%: 0.20±0.011, 0.28±0.02 for top and bottom respectively; Wilcoxon rank sum test, p<0.005). Additionally, and in accordance with previous reports, the activation patches for the corner stimuli showed an anisotropic spread along the anterior-posterior axis (Schuett, Bonhoeffer and Hübener, 2002; Garrett *et al*., 2014; Ji *et al*., 2015b; van Beest *et al*., 2021). In summary, using the corner stimuli we mapped the shape contours onto V1 and found that the point-spread function (PSF) is larger in the anterior part of V1.

### High similarity between the spatio-temporal patterns of neuronal populations evoked by different shapes

Next, we investigated V1 responses to the various shape contour stimuli. Figure 2A shows a sequence of VSD maps evoked by the shapes from an example session. The VSD response shows a large patch of activation in V1 for all stimuli, the black circles denote the corner ROIs of the square shape. All VSD responses showed similar response, initiating from approximately the same region in V1 (Fig. 2A). Notably, despite the clear difference between the large (size: 40×40 deg) shape contour stimuli, the population maps showed highly similar spatial patterns for the different shapes, revealing only subtle differences. To quantify the spatial similarity between the VSD response patterns in the different conditions and across all sessions, we calculate a pixel-wise correlation (r) between each pair of shape response maps (averaged at peak response). The average correlation for all shape pairs was 0.93±0.06 (mean across all pairs; n= 66 which is much higher than the correlation at baseline activity, before stimulus onset (0.47±0.11; Wilcoxon rank-sum, p<0.001).

Figure 2B depicts the TC of the VSD signal in V1 (see Materials and Methods; grand analysis, averaged across all sessions) revealing also high similarity of response dynamics across all shapes (Kruskal-Wallis, p=0.79±0.11 mean±1SEM across shapes and frames). When the visual stimulus was turned-off, the population response decayed towards baseline followed by another transient phase of activation, which is likely an OFF response.

Because the shape stimuli were comprised of thin contours (width 5 deg; i.e. not a filled surfaces), one would expect a lower neural response at the center of the shape contour, as was previously shown in monkeys’ V1 (Zurawel, Shamir and Slovin, 2016; Macknik *et al*., 2019). However, we found the opposite: the population response corresponding to the shape center was higher than the response amplitude at the regions corresponding to the contour edges. To quantify this, we performed a ring ROI analysis (Fig. 2C-E; see Materials and Methods), where the inner ring was positioned at peak response (corresponding also to the mid position between the square corners.) Successive outer ring ROIs were separated by 200µm from each other as shown in Fig. 2C right (example session). The grand analysis for the triangle shape (n=19) confirms that the response amplitude at the center was higher than at the surround (black vs. grays curves, Fig. 2C left). The peak VSD response showed a significant difference between the inner and the most outer ring (Fig. 2D; see also Fig. 2E). This result was reproducible also for the inverted-triangle shape (n=19; inner ring: 1.84×10^-3^±0.16×10^-3^; outer ring 1.10×10^-3^±0.12×10^-3^ Kruskal-Wallis test: p<0.006; with post hoc Tukey test center-surround 1250 µm ring p=0.012; Fig. S1A). Similar differences were found for the square and circle shapes (n=11) however they were n.s. (Square inner ring 1.50×10^-3^±0.24×10^-3^; outer ring 1.02×10^-3^±0.22×10^-3^, Kruskal-Wallis test: n.s. Circle, inner ring: 1.61×10^-3^±0.22×10^-3^; outer ring 1.06×10^-3^±0.19×10^-3^, Kruskal-Wallis test: n.s; Fig. S1A).

To test whether the above results and the high activation at the center emerge from stimulus specific features such as contrast, we performed a similar experiment with a lower contrast (50% contrast; see Materials and Methods, see Fig. S1B). The results were similar to those at high contrast with statistical significance for all shapes (Square, inner ring: 1.58×10^-3^±0.22×10^-3^; outer ring 0.80×10^-3^±0.14×10^-3^ Kruskal-Wallis test: p<0.02; with post hoc Tukey test center-surround 1250 µm ring p=0.04. Triangle, inner ring 1.63×10^- 3^±0.21×10^-3^; outer ring 0.67×10^-3^±0.15×10^-3^, Kruskal-Wallis test: p<0.004; with post hoc Tukey test center-surround 1250 µm ring p=0.007. Inverted-triangle, inner ring: 1.78×10^- 3^±0.26×10^-3^; outer ring 0.71×10^-3^±0.15×10^-3^, Kruskal-Wallis test: p<0.01; with post hoc Tukey test center-surround 1250 µm ring p=0.02).

In summary, different large shapes evoked similar activation patterns with peak VSD response at the center of the shape, an observation that was reproducible for different contrasts. We next aimed to investigate a possible explanation for the high response at the center of the shape.

### Predicted activation patterns based on point-spread function explain the high activation at the center of the shape

A possible explanation for the high activation at the response center can be related to the size of the cortical PSF, defined as cortical area evoked by a point stimulus (Grinvald *et al*., 1994; Frostig *et al*., 2017). Notably, the PSF as measured with VSDI extends over a large cortical area because it reflects both the sub-threshold and supra-threshold membrane potentials (i.e. spiking activity) of neuronal populations.

To study if the peak VSD response at the center can be explained by the PSF, we performed the following analysis (see Materials and Methods, Fig. 3). First, we normalized the VSD map evoked by each corner stimulus (Fig. 3Ai) and fitted a 2D Gaussian model to the activation pattern (Fig. 3Aii, PSF kernel). Next, we convolved the shape stimulus projection on V1 (Fig. 3Aiii) with the PSF kernel to compute the expected pattern of activation (Fig. 3Aiv). We used two approaches to compute the expected VSD response for each shape: (1) the stimulus projection was convolved with a single PSF (Fig. 3A) (2) each corner (and neighboring edges) of the projected stimulus were convolved with the same corner’s PSF (for example top corner convolved with the top triangle stimulus part). The second approached was used in order to account for the difference in anterior and posterior PSF (Fig. 1E).

To evaluate the similarity between the expected and measured shape pattern, we calculated the pixel-wise correlation (r) for each pair of observed and predicted maps (Fig. 3B). We found high correlation for all the stimuli (triangle: range 0.77±0.02 to 0.80±0.03; inverted-triangle: range 0.70±0.033 to 0.73±0.035; square: range 0.66±0.06 to 0.67±0.06; circle: range 0.73±0.034 to 0.75±0.036), meaning high similarity between the observed and predicted maps. Finally, Figure 3C shows for the predicted maps, that the response at the center is higher than outer region (Kruskal-Wallis test: p<0.0001) similar to the stimuli evoked maps (Fig. 2C-D). This result was reproducible for all predicted shape maps (Fig. S2).

In summary, the results suggest that the spatial patterns evoked by the contour stimuli and the high activation at the center can be explained by the PSF.

### Narrow and wide parts of the shape contour generate wide and narrow spatial profile of cortical response

Previous behavioral studies reported that mice can discriminate between shapes (Bussey, Saksida and Rothblat, 2001; Brigman *et al*., 2005). Thus, despite the high temporal and spatial similarity of the activation patterns evoked by the various shapes (Fig. 2A-B, Fig. S1A), one would expect for some differences in the neuronal activation patterns. Based on the retinotopic map in V1, it is expected that the width of spatial profile of cortical activation will co-vary with the local shape width. For example, the spatial profile of cortical activation evoked by the narrow apex of the triangle is likely to be narrower than the spatial profile of cortical activation evoked by the wide triangle base. Thus, our next step was to quantify the spatial profile of cortical activation for the local width variation of each shape. For this purpose, we computed three different spatial cuts across the anterior-posterior axis of V1 corresponding to the triangle base and apex (Fig. 4A and see Materials and Methods). Next, we computed the spatial profile (by averaging over the width of each spatial cut) as function of time to obtain the space-time maps. Figure 4A bottom shows the space-time maps, depicted separately for each spatial cut (averaged across all animals). The space-time maps (zero on the y-axis denotes peak response) show a narrower spatial spread of activation in the posterior cut (Fig. 4A, right panel) as compared with the anterior cut (Fig. 4A, left panel). This result is expected: a narrower profile of cortical activation corresponding to the narrow apex of the triangle as compared with the wide base of the triangle.

Next, we aimed to quantify the width of the spatial profiles (Fig. 4B and see Material and Methods; width is normalized to peak response). As shown in Figure 4A, we assumed that the width of the spatial profile should co-vary with the narrow apex and wide base of the triangles. For the triangle shape (Fig. 4B top left), the width of the anterior spatial profile (corresponding to the triangle base), was larger (n.s.) than the width of the posterior profile (corresponding to the triangle apex). For the inverted-triangle (Fig. 4B top right) opposite results were observed (n.s.). However, the spatial profile of the symmetrical shapes (square and circle, Fig. 4B bottom) showed a larger width for the anterior spatial cuts (compared with the posterior cuts) that reflects the larger PSF of the anterior part (Fig. 1E).

Thus, to account for this feature in the retinotopic map, we defined a difference index (see Materials and Methods Eq. 1; Fig. 4C) which is a normalized measure for the spatial profile width of a test shape, relative to the symmetric square shape. Figure 4C shows the difference index for the triangle, inverted-triangle and circle for the 3 cuts. In the posterior cut (corresponding to the upper VF; Fig. 4C right) the difference index showed a higher value for the triangle (0.07±0.02; p<0.005 sign-rank test from zero) than the inverted triangle (0.05±0.02; n.s. difference from zero, sign-rank test). These results indicated that the spatial profile of the square is larger than the apex of the triangle but similar to the base of the inverted-triangle. In the anterior cut, opposite results were obtained: the inversed triangle had a large value (0.10±0.03; p<0.05 sign-rank; triangle: 0.04±0.01; n.s. difference from zero, sign-rank test). Interestingly, also in the middle cut, the difference index for both triangle and inverted-triangle showed a larger and significant value, meaning that the square stimulus generated a wider spatial profile, as expected (triangle: 0.08±0.02; p<0.005 sign-rank test; inverted-triangle: 0.12±0.02; p<0.0001 sign-rank test). In addition, the difference index for the triangle stimuli was larger than the expected threshold (dashed line, Fig. 4C; see Materials and Methods). Finally, for the circle shape, all 3 spatial cuts showed a difference index that is around zero (from anterior to posterior (mean ± 1SEM: A: 0.02±0.03; M: 0.01±0.02; P: 0.04±0.03; n.s. difference from zero, sign-rank test). This means similar activation width for the square and circle stimuli.

In summary, these results suggest that the local variation of stimulus width correlates with spatial profile of cortical activation. In other words, the narrow and wide parts of the triangle shape contour generate wide and narrow spatial profile of cortical response.

### Differential maps of stimulus-pairs reveal informative pixels for shape discrimination

Our next step was to investigate which pixels in V1 hold information for discriminating between the shapes. For this, we computed a pixel-by-pixel differential activation map for each pair of shapes (see Material and Methods and Fig. 5A) and then calculated the mean value of V1 pixels in the differential map, and set a threshold of Mean±1STD. Next, we created a STD map by defining three classes of pixels: (i) pixels with values smaller than Mean-1STD (-1, brown color; Fig. 5A). (ii) pixels with values larger than Mean+1STD (+1, yellow color; Fig. 5A). (iii) pixels with values within Mean±1 STD (0, orange color). Pixels with values equal or larger than ±1 denote informative pixels for shape discrimination (see Material and Methods).

Examples of the STD maps for triangle and inverted-triangle, from two animals, appear in Figure 5A. Notably, the maps show that during stimulus presentation there are clear negative and positive pixel patches (brown and yellow pixels, respectively; Fig. 5A right maps for each animal) whereas during baseline (before stimulus presentation) there are no clear patches (Fig. 5A left maps for each animal). Positive and negative pixel patches are expected for positive and negative activation difference for the shapes pair. For the triangle and inverted-triangle pair, the positive and negative patches appear, as expected from the retinotopic organization, in the anterior (lower VF, corresponding to the triangle base) and posterior (upper VF, corresponding to the triangle apex) parts of V1, accordingly (see illustration in Fig. 5A top). The blue points represent the center-of mass (see Materials and Methods) for the positive or negative patches and they appear at close proximity to the bottom and top corner ROIs, accordingly.

To quantify the relation between the patches’ location and the shape projection onto V1 we computed the distance between each patch location (i.e. center-of-mass) and the corner ROIs. The distance results are depicted in Figure 5B left panels. For the triangles pair (Fig. 5Bi) we found that the negative patches were closer to the top corner ROIs (0.41±0.03) than to the bottom corner ROIs (0.86±0.05; p=1.2×10^-4^ rank-sum test). This means that in the posterior part of V1, the triangle apex evoked a smaller activation than the base of the inverted-triangle evoked (see illustration in Fig. 5A top). For the positive patch, opposite results were obtained, meaning that triangle base evoked higher activation in anterior part of V1 relative to the apex of the inverted triangle (top: 0.89±0.05; bottom: 0.53±0.06; p=3.6×10^-6^ rank-sum test).

For the triangle and square pair (Fig. 5Bii), the negative patch was closer to the top corners (top: 0.58±0.15; bottom: 0.86±0.06; p=0.05 rank-sum test), meaning that the top square edge evoked higher activation than the triangle apex. Interestingly, the distance of the top or bottom corners from the positive patches in this shape pair, was similar (n.s. different), suggesting that the position of the positive patches was not related to specific corner ROIs. Similar results was obtained for the square and circle shapes (Fig. 5Biii), that have a large spatial overlap: the distance of the positive and negative patches from the top or bottom corners were not significantly different (positive: top 0.77±0.20; bottom 0.90±0.07 n.s.; negative: top 0.81±0.18; bottom 0.67±0.09 n.s.). Same results were obtained for the scrambled triangles (Fig. 5Biv) as one could expect from the spatial overlap (positive: top 0.93±0.09; bottom 0.90±0.12; n.s.; negative: top 0.68±0.14; bottom 0.64±0.22; n.s.)

As we showed earlier, the peak of response activation for all shapes, occurred at the center of the evoked pattern (Fig. 2C-E, Fig. S1A, Fig. 3C, Fig. S2). Thus, we asked: can the response at the center hold information for discrimination between the shapes? To answer this question, we calculated the distance of each patch from the center. For all shape-pairs we found similar distances between the center and the negative and positive patches (Fig. 5B; n.s. Wilcoxon rank-sum test), meaning that the pixels in the center did not hold information for shape discrimination.

To further quantify whether the cortical regions around corner ROIs show high or low values in the STD maps, we averaged the values (-1, 0 or 1) in a circle ROI aligned on the top or bottom corners (the two top corners and the two bottom corners showed similar result of STD values and thus where averaged; see also Fig. S3B). Figure 5Bi right show the results for the triangles-pair: the average STD map for the top corner ROI show mainly red-brown pixels, i.e. negative STD values (mean=-0.23; posterior area). The average STD map for the bottom corner ROI shows a yellow color i.e. positive values (mean=0.17; anterior area), as expected from the retinotopic organization (see illustration in Fig. 5A top). The histogram below the maps depict the distribution of the average STD values (arrows indicate the mean). For the triangle pair the two histograms are well separated showing only a small overlap. The averaged STD map for the square-circle pair and scrambled pair (left panels in Fig. 5Biii, iv) show mainly orange color i.e. values close to zero (i.e. pixels that are not informative for discrimination). The histograms below the maps show the two distributions with high overlap between the top and bottom distribution. In addition, both histograms are centered around zero, meaning, not significant change between the shapes. Finally, we note that similar results were obtained for a lower stimulus contrast (50%; see Materials and Methods and Fig. S3A) further demonstrating that the results are not specific to 100% contrast.

In summary, the above results suggest that V1 in mice encode shape discrimination based on low level cues i.e. local shape differences of the stimulus.

### Comparison of shape processing in monkey’s V1 foveal representation and mouse V1

Mice are an important animal model to study the visual system, yet, there are some key differences between their visual system and that of primates. Humans and NHPs use their fovea to inspect objects with high resolution whereas mice lack both fine-scale visual acuity and the specialized fovea region in the retina (Huberman and Niell, 2011). Thus, our next step was to compare shape processing in V1 of mice and monkeys while focusing on the foveal representation of the monkey’s V1. For this purpose, a fixating monkey was presented with various small (0.8 deg) shape contours stimuli: circle, rectangle, and triangle at the fovea area (Fig. 6A left; see Materials and Methods). Figure 6A middle shows the VSD maps for the shape stimuli, from an example session. The spatial patterns of population responses to the shape contours that appear in the monkey’s fovea, show clear resemblance to the presented shape contour with large differences between the shapes (Fig. 6A left). We note that in a recent work, we showed that the VSD activation maps for the shape stimuli can be well predicted from a 2D analytical model of a retinotopic map convolved with a PSF (see Zurawel, Shamir and Slovin, 2016). In contrast, the evoked VSD response in mice V1 (Fig. 6A right) showed a large patch of activation w/o a clear similarity to the visual stimulus and high similarity among the stimuli.

To quantify the spatial resemblance between the VSD response patterns of the different shapes, we calculate the pixel-wise correlation (r) between each pair of shape response maps. Interestingly, while in mice the correlation was high (0.93±0.06 for all shape pairs) in the monkey, we found a significantly lower correlation (0.62±0.02; rank-sum test p<0.001 for all shape pairs; correlation at baseline in monkey is: 0.057±0.04). This result may suggest a topographic code in V1 fovea of monkeys while mice may need to use local feature difference i.e. low level cues for discrimination.

To quantify the spatial patterns in monkey and compare it to mice, we performed a ring analysis on the VSD maps in monkeys (as in Fig. 2C-E for mice). For this, we generated a continuous set of non-overlapping rings (Fig. 6B right; see Materials and Methods). Interestingly, the VSD TC averaged over each elliptical ring ROI showed an opposite result to that in mice: the inner ring showed a lower activation while the outer most ring showed a higher activation, for circle stimulus. This is further quantified in Figure 6C that shows the peak response amplitude for each ring. A gradual increase in amplitude can be seen, with the lowest amplitude at the center region of the pattern (from the inner ring: 7.06×10^-4^±0.60×10^-4^ to the outer ring 9.7×10^-4^±0.67×10^-4^, mean±1SEM. Kruskal-Wallis test p=0.0005). Figure 6D shows a 3D map of the evoked response averaged across all circle sessions. The center of the 3D map shows a clear lower activation, i.e. hole at the center of the activation (Zurawel, Shamir and Slovin, 2016; Macknik *et al*., 2019), whereas the VSD response at the circle contour is higher. In summary, shape processing in the foveal region of a monkey’s V1 and in mice V1 shows a clear different processing for shape contour stimuli. Suggesting that population response in V1 of a monkey shows a high resolution topographic code whereas in mice the VSD map suggested blurred representation of the shape.

### Visual stimuli at increasing eccentricity from the fovea in monkeys: the VSD response pattern in a parafoveal region is more similar to that in mice V1

The mouse retina is considered as a model for peripheral vision (> 6-7 deg) in primates. Although in the V1 imaged area in our monkeys we did not have access to these peripheral eccentricities of the VF, we did have access to more parafoveal locations. Thus, in another set of experiments we analyzed shape responses from both foveal and parafoveal regions of monkey’s V1 (Métin, Godement and Imbert, 1988; Jeon, Strettoi and Masland, 1998a). The monkey was presented with a single circle contour stimulus at three eccentricities ranging from parafoveal to foveal locations. The VSD responses are shown in Figure 7A left (an example session) and demonstrate that at the largest eccentricity the population response pattern appears as a small elliptical blob (Fig. 7A top). This pattern transforms into a large annulus with a hole at V1 foveal region (Fig. 7A bottom). To quantify this, we set a spatial cut on the VSD maps for each stimulus location and computed a space-time map as demonstrated in Figure 7A right (see Material and Methods). The space-time map for the largest eccentricity (Fig. 7A, top-right) shows one peak at the center of response (zero in the y-axis), whereas, in fovea (top panel) two peaks are observed with low activation at the center of the shape (see Materials and Methods). To compare this with the mouse responses, we set a spatial cut over V1 activation in the mouse (Fig. 7B left) and computed the space-time map (Fig. 7B right). Interestingly, the population response in mouse V1 show a spatial pattern that reminds the largest eccentricity in the monkey’s data: one peak at the center. Figure 7C shows the normalized spatial profile (at peak response; see Materials and Methods) for monkey (black and grays curves) and mice (red curve). The parafovea profile (light gray curve) shows a similar pattern to the spatial profile in the mouse (red curve) with one peak at the center (distance=0, x-axis). However, the fovea (black curve) and the near parafovea (dark gray) spatial profiles show 2 peaks with low activation at the center. To quantify the differences between the cortical spatial profiles of mice and monkey responses, we calculated a center-edge index (see Materials and Methods; Fig. 7D). A positive value indicates higher response at the center relative to edge and a negative value means higher response at the edge vs. the center. As shown in Figure 7D mice (red) and most parafovea location (light gray) show positive values, meaning higher response at the center of the spatial cut, with similar center-edge value (n.s. rank-sum test; significant from zero p<0.01 sign-rank test). Whereas, negative values obtained for fovea and close to fovea location and were significant different from mice results (rank-sum test p<0.001 with Bonferroni correction; significant from zero sign-rank test p<0.001). These results suggest a larger similarity between visual processing in mice V1 and in parafoveal areas of the monkey, as compared with foveal regions in monkey.

### Seed correlation analysis reveals the shape contours in monkey

Synchronization is an important neural mechanism for object encoding. To study the neural correlation involved in shape processing, we computed synchronization maps at the single trial level (see Materials and Methods). Using a sliding window of 100 ms we calculated the Pearson correlation coefficient (r) for all pixel pairs between seed ROI pixels and all other pixels within the imaged visual cortex (Eq. 4; Fig. 8A). This was done after removing the direct stimulus contribution by subtracting the mean (across trials) visually evoked response from each pixel (see Eq. 3 in Materials and Methods). Next, the correlation maps were averaged across trials and for all seed pixels, resulting in synchronization maps depicted in Figure 8B as time sequence maps or time-averaged maps as in Figure 8C (top for mouse and bottom for monkey). The synchronization maps show the zero-time lag correlation between the seed pixels and the pixels in all the imaged area, during visual stimulus presentation (Maatuf, Stern and Slovin, 2016; Margalit *et al*., 2022). We further analyzed only pixels with correlation exceeding the shuffle values (see Materials and Methods). We note that the correlation maps reveal the spatial synchronization pattern in V1 area (that is no related to the direct contribution of the visual stimulus, because the latter was subtracted from the data).

Figure 8B-C top shows an example for synchronization maps in mouse where the seed ROI is marked with a red circle. A patch of higher correlation (white pixels) appears in V1, especially at pixels surrounding the seed ROI. Figure 8C bottom shows an example for synchronization map in the monkey (seed ROI is denoted by gray circle). Remarkably, in monkey the synchronization pattern is elliptical, reminiscent to the neural pattern evoked by the circle stimulus. To quantify the synchronization in V1 we calculated the mean correlation between the seed ROI pixels to others ROI pixels (as defined below) across all sessions. We defined 3 types of ROIs in V1 (color coding of the ROIs appears in Fig. 8Eii) as shown in Figure 8D top (see also Materials and Methods): (1) *Related ROIs*: shared corner ROIs, i.e. ROIs with overlapping stimulus input for the shape-pair. (2) *Un-related ROIs*: corner ROIs with non-overlapping stimulus input for the shape-pair. (3) *Center ROI*: located at the center of the shape contour. Next, we calculated the correlation between the different ROIs as depicted in Figure 8D bottom (see Materials and Methods). *(1) Related correlation*: the correlation between a seed related ROI to all other related ROIs; *(2) Un-related correlation*: the correlation between seed related ROI to all other un-related ROIs. *(3)) Center correlation*: the correlation between center ROI to all seed related ROIs.

Figure 8Ei shows the related, un-related and center correlations in mice, for all shape contour stimuli (n=19 for triangles; n=11 for square and circle). We hypothesized that neuronal population whose RF are falling on the shape contour, i.e. related correlation, will show higher correlation than neuronal population whose RF are falling elsewhere. Indeed, the related correlations were significantly higher than the un-related correlation for all shapes (p<0.01 or p<0.001, two-sided Wilcoxon rank sum test, Bonferroni correction). This finding is consistent with the role of synchrony for encoding stimulus features belonging to the same object. Surprisingly we found that the center-correlation showed the highest correlation (although there is no direct stimulus input at the shape’s center), higher than the unrelated correlation (p <0.001, for all shapes) and related correlation (n.s., except for circle p<0.05). This finding may suggest that in mice, the center region was correlated with all other shape parts, that may lead to visual perception of a blurred pattern.

Interestingly, the pattern of correlation was different in the monkey. Figure 8Eii shows that the highest correlation appeared at the related ROIs. Interestingly, the correlation for the center ROI showed a lower value whereas the lowest correlation was found for the un-related seed ROIs. This pattern was reproducible for all shapes. Finally, to compare between mice and monkey correlation results, we computed a correlation modulation index (CMI, Fig. 8F; see Materials and Methods Eq. 5) between different categories, indicating which correlation was higher. The correlation between related vs. center showed negative CMI values for mice (red), but for monkey (black) the CMI was positive, meaning higher center correlation than related correlation in mice and an opposite result for monkey. The CMI value for related vs. un-related were positive for both animals with higher CMI for monkey. The CMI values for the un-related vs. center correlations were negative for both animals, with much larger values for monkey (all CMI where significant from zero; sign-rank test p<0.001). In summary, the CMI results suggest that synchronization patterns are different for mice and monkey.

## Discussion

We investigated shape processing in mouse V1 and compared it to population responses evoked by shape stimuli in the foveal region of monkey’s V1. The activity in mice was characterized by a wide spread activation with peak response at the shape’s center and small visible differences between shapes. In contrast, the evoked spatial patterns of neural activation in monkey’s V1 resembled roughly the shape of the original stimuli with a clear hole at the center. The foveal representation in V1 of primates demonstrated clear neural differences at sub-degree resolution of the shapes. Thus, V1 activity in both species can discriminate between different shapes but may use a different neural strategy.

### Non-homogeneous point spread function in the retinotopic organization of V1

A retinotopic map is a key feature of V1 in humans, monkeys (Tootell *et al*., 1988; Blasdel and Campbell, 2001) and rodents (Schuett, Bonhoeffer and Hübener, 2002; Huberman and Niell, 2011; Garrett *et al*., 2014; Macknik *et al*., 2019). A high-resolution retinotopic map is an essential requirement for neural encoding of objects at fine details and the ability to discriminate between shapes. Using corner stimuli we mapped the shapes onto mice V1 and found as in previous studies, that the upper and lower VFs are projected onto the posterior and anterior parts of V1 (Fig. 1A-C; Schuett, Bonhoeffer and Hübener, 2002; Wang and Burkhalter, 2007; Ji *et al*., 2015a; Zhuang *et al*., 2017).

Past studies reported on a homogeneous representation of the VF in mouse V1 and a nearly uniform cortical magnification factor (CMF) without a clear foveal area which holds a significant topographic variation in primates (Jeon, Strettoi and Masland, 1998b). Yet, recent papers reported on non-homogeneities in mouse retina that may be beneficial for behavioral purposes. Retinal ganglion cells (RGC) subtypes display distinct topographies in the mouse retina (i.e. density and cell morphology), suggesting that the different subtypes serve to encode visual features encountered in specific regions of the VF to optimally sample the visual environment (Warwick *et al*., 2018; Heukamp, Warwick and Rivlin-Etzion, 2020). Another study reported on a slight increase in CMF at the binocular zone of mice V1 (Garrett *et al*., 2014) while van Beest et al. (2021) suggested that V1 contains a region of enhanced spatial resolution (a ’fovea-like’ region) in the binocular upper VF (van Beest *et al*., 2021).

Our results are consisted with an inhomogeneous visual representation in mouse V1. Interestingly, we found that the PSF was larger in the anterior part of V1 (i.e. the lower VF) than in the posterior part of V1 (i.e. the upper VF, Fig. 1E). In addition, and in accordance with previous studies, we found anisotropy of the visual responses along the anterior-posterior axis (Schuett, Bonhoeffer and Hübener, 2002; Garrett *et al*., 2014; Ji *et al*., 2015b; van Beest *et al*., 2021). How can inhomogeneous cortical representation meet the ecological requirements of mice living in natural enviroment? Different regions in V1 receive information from different parts of the VF, and thus may encode visual information that is behaviorally relevant to those parts of the VF. For example, the upper VF is mainly concerned with predator detection whereas the lower VF, holds visual information for foraging in mice (Singleton and Krebs, 2007; Heukamp, Warwick and Rivlin-Etzion, 2020). These different visual functions may utilize different neural mechanism and the large PSF at the lower VF may facilitate object processing, an essential capability for foraging.

### High activation at the center of the contour in mice but not in monkey’s fovea

A shape contour stimulus in the mouse, evoked peak response activation at the center of the shape (Fig. 2C-E), despite the hole at the shape’s center, whereas in monkeys, a shape contour in V1 foveal region evoked activation pattern with a clear hole at the center of the shape (Fig. 6). These different patterns raise a question whether the same stimulus is processed and perceived in a different manner in these two animal species.

We showed that in mouse the high activation at the shape center can be explained by convolving the PSF with the stimulus projection in V1 (Fig. 3). The predicted maps showed higher activation at the center, similar to the observed pattern evoked by the shape contour (Fig. 3C). The large PSF in V1 of mice related to the RFs that are much larger (Métin, Godement and Imbert, 1988; Niell and Stryker, 2008) than the RFs in monkeys, particularly in the foveal area (0.1 deg; Wang and Burkhalter, 2007; Murgas et al., 2019). The large size of RFs combined with their spatial overlap can lead to large spatial summation of neuronal activation in the mouse. We note that estimation of the PSF and its effects on the encoding model of a shape contour stimulus in monkeys was previously shown in Zurawel et al. (2014 and 2016).

The peak response at the center of the shape contour in mice raises an interesting question: what is the emerging visual perception from this pattern, a blurred surface or a fine contour? How can a mouse in the wild discriminate between a dark surface i.e. a hole in the ground from a contour? In different set of experiments, mice were presented with a luminance surfaces (i.e. white or black large filled squares). Interestingly, we found that the TC of the VSD signal evoked by a surface or contour at the shape center was different (data not shown). This suggests a different neural code for surfaces and contours and thus possible discrimination between these two stimulus types. Future behavioral studies should examine the capability of mice to discriminate between surfaces and contours.

### Mice visual system can discriminate between shape contour

Previous behavioral studies reported that the mice can perform complex visual tasks such as: figure-ground segregation (Schnabel *et al*., 2018), object and shape recognition (Bussey, Saksida and Rothblat, 2001; Brigman *et al*., 2005) and discrimination between natural images (Yu *et al*., 2018). These observations argue for the existence of cortical circuitry to support these capabilities. We found that different large shape stimuli evoked wide spread neural activation in V1 of mice with high spatio-temporal similarity (Fig. 2A, 6A). We further showed that although mice have low visual acuity, the spatial profile of V1 activation co-varied with the local shape width (Fig. 4). Consisted with this result, the STD maps of shape-pairs showed negative and positive patches at non-overlapping regions of the shapes, meaning that in specific V1 regions, one shape evoked a lower or higher activation from the other (Fig. 5). Our findings support the behavioral notion that mice can discriminate between shapes based on local cues, i.e. local differences of the shapes that are reflected in the neural responses. These findings were reproducible for shape stimuli at different contrast levels (Fig. S3).

### Shape processing: holistic approach versus low-level cues

V1 in mice show hallmark features that were previously reported in primates, including tuning curves to orientation (Dräger, 1975; Niell and Stryker, 2008; Glickfeld, Histed and Maunsell, 2013) and temporal or spatial frequency (Marshel *et al*., 2011; Palagina, Meyer and Smirnakis, 2017; Marques *et al*., 2018) as well as sensitivity to contrast and luminance changes (Dai and Wang, 2012). However, key functional maps such as ocular dominance or orientation columns, which are well established in higher mammals, were not found in the mouse visual cortex (Blasdel and Salama, 1986; Ts’o *et al*., 1990; Ohki and Reid, 2007; Van Hooser, 2007). Moreover, while primate use their fovea to inspect objects at high details, mice lack both fine-scale visual acuity and the specialized fovea region in the retina (Prusky, West and Douglas, 2000; Van Hooser, 2007; Huberman and Niell, 2011). Thus, it remains unclear to what extent the two species share a common representations of visual objects. In particular, it is unclear whether mice utilize low-level features to perform shapes discrimination or they apply a holistic strategy?

Primates’ ventral stream carries information related to objects form and recognition and is invariant to object position or size. This transformation tolerance (Ratan Murty and Arun, 2017b) indicates that monkeys process shape in a holistic manner. A number of behavioral studies suggested the existence of a similar holistic approach in rats (Zoccolan *et al*., 2009; Tafazoli, di Filippo and Zoccolan, 2012; Vermaercke and Op De Beeck, 2012; Alemi-Neissi, Rosselli and Zoccolan, 2013; Vinken, Vermaercke and Op de Beeck, 2014). However, Luongo et al. (2021), investigated in mouse the neuronal mechanisms underlying figure-ground segmentation, and compared it to other species (Luongo *et al*., 2021). They found that mice and primates use fundamentally different strategies beyond visual resolution. Unlike primates, mice are strong texture-dependence, suggesting that they do not process shape stimuli in a holistic manner but rather use low-level cues. Interestingly, the authors trained mice on relevant behavioral tasks and showed that mice are incapable of texture-invariant segmentation. These results add to a growing body of evidence showing that mice use low-level cues to discriminate between different shapes (Minini and Jeffery, 2006; Yu *et al*., 2018). Yu et al. (2018) examined if mouse can discriminate between natural images, and found that mice attended to specific sub-regions of the image and also to the edges orientation within that sub-region in order to discriminate between images. Furthermore, Minini and Jeffery (2006) found that rats relied on local luminance differences especially at the lower hemifield (Minini and Jeffery, 2006). This is consistent with our results and with previous studies (Lashley, 1938; Simpson and Gaffan, 1999).

Interestingly, Luongo et al. (2021) suggested that the strong texture-dependence in mouse segmentation behavior suggests a visual strategy more similar to deep networks than primates do (see also You and Mysore, 2020). Recent work has shown that deep networks, unlike humans, fail to exploit global image features such as object boundaries and shape to perform classification, but rather, strongly rely on texture features (Van Nes and Bouman, 1967; De Valois, Morgan and Snodderly, 1974). Our findings are consistent with the argument that mice use low-level cues to discriminate between shapes. We show that the differences between shapes occur in local regions within V1, where the shape projections on the retinotopic map in V1 do not overlap (see Fig. 5A).

### The response in mice V1 is more similar to monkey’s parafoveal region

In primates, visual acuity is highest in the fovea and it decreases with eccentricity. This feature results from several changes along the eccentricity of the retina (Bringmann *et al*., 2018): (1) The density of RGCs is decreased and their dendritic arbor size increases with eccentricity (Field *et al*., 2010). (2) The pooling of information from retinal receptors by RGC is greater in periphery than in fovea (Curcio and Allen, 1990). (3) The fovea is comprised from cones and avoid rods, however in peripheral retina there is an opposite pattern, and the rods outnumbers the cones (Curcio *et al*., 1990). (4) The RFs size in primates’ fovea are small and increase with eccentricity (Keliris *et al*., 2019). Finally, in V1 the CMF decreases with eccentricity, meaning that information must be enlarged to be solved in the parafovea area (Strasburger, Rentschler and Jüttner, 2011). The differences between fovea and more peripheral vision may imply that the visual processing in these areas have different computational capabilities derived by distinct neuronal circuit.

Several studies found that visual performance and spatial resolution are better in the fovea than in parafovea (Pointer and Hess, 1989; Decaux and Soupart, 2001). It was found that as a visual target shifted from foveal locations towards more peripheral locations, the observers’ performance decreased and their reaction time increased (Carrasco *et al*., 1995). Furthermore, visual capacity for letter stimuli decreases with eccentricity (Staugaard, Petersen and Vangkilde, 2016) and the ability to process emotional facial expressions (Bayle *et al*., 2011; Guo, Liu and Roebuck, 2011) decreases with eccentricity.

Interestingly, the mouse entire retina is more similar to the peripheral retina of primate rather than fovea area. It was found that the mouse retina is rods dominated with uniform distribution across all retina (Jeon, Strettoi and Masland, 1998a), similar to primates peripheral region. Furthermore, the cone density is similar between mice and primate eye at 6-7 deg away from the fovea (Jeon, Strettoi and Masland, 1998a). Moreover, the RFs in mice are relatively large ∼14 deg (Métin, Godement and Imbert, 1988), similar to RFs at the monkey’s periphery. Although we did not have access to cortical peripheral processing, we aimed to study how shape contour is represented in fovea and parafovea areas and whether the neural representation in the parafovea area will be more similar to mice. Thus, we investigated the VSD responses to a circle shape at increasing eccentricities: from fovea to parafoveal locations and compared it to the VSD response in mice (Fig. 7). In the fovea, the VSD response to a small circle stimulus resembled roughly the shape of the original stimuli, i.e. showing a topographic code. Interestingly, the VSD evoked spatial pattern became blurred and more similar to the mice at the largest eccentricity, where both animals show one peak at the center of the response. These results, support the differences between fovea and parafovea vision and may imply that the visual processing in these areas have distinct neuronal circuits. Moreover, the similarities between mice and primates parafovea may suggest that visual capabilities in mice are more similar to the those of monkey’s parafoveal regions.

### Seed correlation analysis revealed different shape processing in mice and monkey

Synchrony was suggested as an important neural mechanisms for feature integration of an object (Gray, 1999; Singer, 1999). It was suggested that neurons representing the visual object are synchronized in their activity (binding theory), and therefore are distinguished from other neurons that encode other parts of the visual scene (Eckhorn *et al*., 1988; Gray *et al*., 1989; Singer and Gray, 1995; Kreiter and Singer, 1996; Castelo-Branco *et al*., 2000; Gail, Brinksmeyer and Eckhorn, 2000).

We investigated synchrony in mice V1 area for shape stimuli by computing synchronization maps where the seed pixels were located on the shape edges (i.e. corner ROI; Fig. 8A). We hypothesized that neurons whose RF are falling on the shape contour (i.e. the corners ROIs) will show higher correlation than neurons whose RF are falling elsewhere. Indeed, we found higher correlation between regions in V1 that overlap with the shape (’related correlation’). In contrast, un-related regions in V1 that had no-overlap with the shape showed lower correlation (’un-related correlation’). These results are in accordance with the binding or synchronization hypothesize. Interestingly, although we used stimuli comprised of shape contour, the correlation between the region representing the center to all other regions, i.e. the ’center correlation’ was highest in mice, even larger than the ’related correlation’ (Fig. 8E). This finding may suggest that in mice, the center area was correlated with all other shape parts (Fig. 2D-E), further suggesting visual perception of a blurred pattern. Interestingly, in monkey we found that the ’related correlation’ is larger than the ’un-related correlation’, as shown for mice. However, in contrast to the mice, the correlation at the center was lower than the ’related correlation’ (Fig. 8E) and this was further quantified by the modulation index quantifies. These results offer further support to the notion that mice and monkeys use different neural code to discriminate between shapes.

