## Supplementary material for "Different patterns of visual processing for shapes in V1 of mouse and monkey"

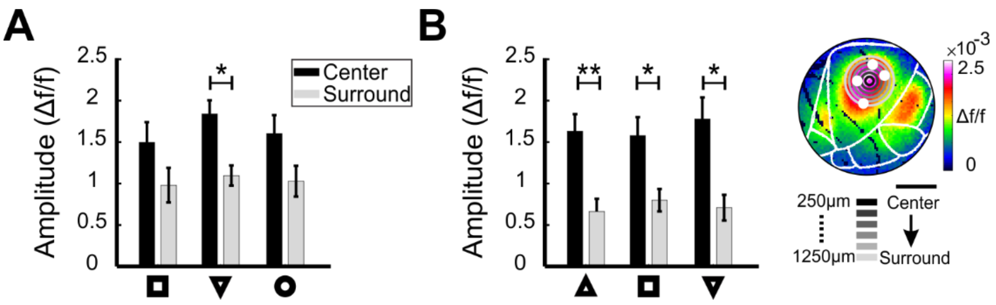


***Supplementary Figure S1*. Ring analysis for 100% and 50% contrast in all contour shape stimuli**

(A) Peak response (time averaged 150-300 ms post stimulus onset) for all sessions in center and surround ring ROIs, for all shapes on black background. Kruskal-Wallis test: p=0.006, with *post hoc* Tukey tests: p=0.012, for inverted triangle. (B) Right: Schematic illustration of ring ROIs superimposed on the VSD map evoked by the triangle shape on gray background. The distance between two successive rings is 200 μm. White filled circles denote the corners ROIs. Left: Peak response (time averaged 250-400 ms post stimulus onset) for all sessions in center and surround ring ROIs, for all shapes on gray background. Kruskal-Wallis test: p=0.004, p=0.04, p=0.01 with *post hoc* Tukey tests: p=0.007, p=0.04, p=0.02, for triangle, square and inverted triangle, respectively).

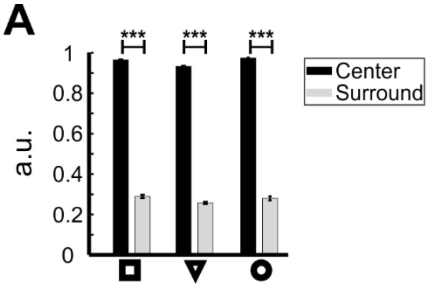


***Supplementary Figure S2.*** **Ring analysis for all predicted contour shape stimuli**

Ring analysis as in Fig. 3C but for the predicted response of all stimuli (Kruskal-Wallis test: $p<0.0001$ with *post hoc* Tukey tests: center ring with surround ring $p<0.001$).

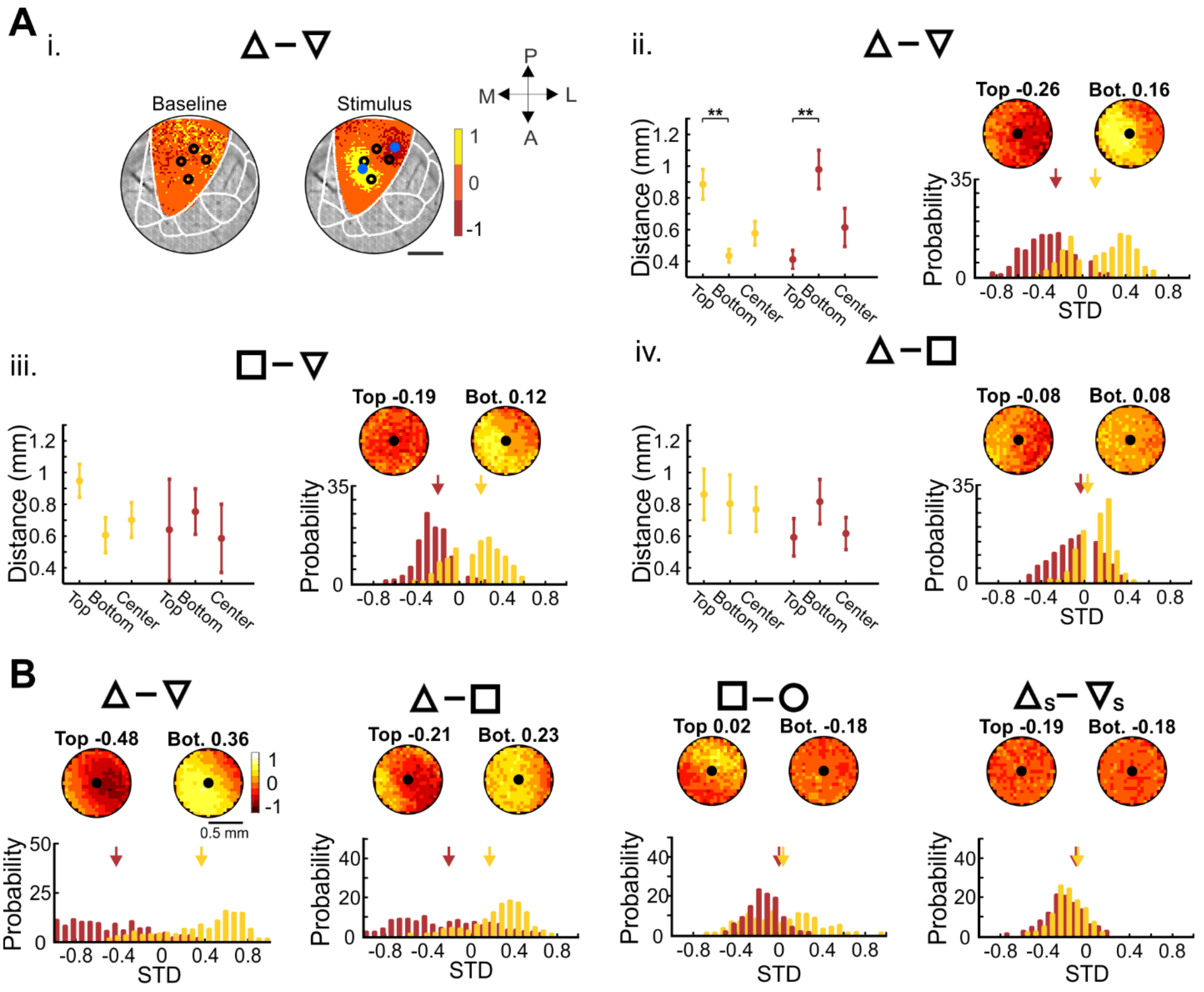


***Supplementary Figure S3.*** **Differential STD maps for stimuli pairs presented over gray background and STD maps for contour shape stimuli in 100% contrast**

**(Ai)** An example of STD maps for a triangles-pair (triangle and inverted-triangle) presented over a gray background, 50% contrast. Same as Figure 5A. **(Aii-iv)** same as Figure 5B but for gray background (50% contrast; n=6, map radius is 0.5 mm). Wilcoxon rank-sum tests: ** $p<0.01$. **(B)** same as Figure 5B right but for non-binned STD maps, i.e. the mean value of the differential map was subtracted from each pixel and then divided by the STD value of the map.
